# Insights into the drivers of radiating diversification in biodiversity hotspots using *Saussurea* (Asteraceae) as a case

**DOI:** 10.1101/2021.03.15.435394

**Authors:** Xu Zhang, Jacob B. Landis, Yanxia Sun, Huajie Zhang, Tao Feng, Nan Lin, Bashir B. Tiamiyu, Xianhan Huang, Tao Deng, Hengchang Wang, Hang Sun

**Affiliations:** CAS Key Laboratory of Plant Germplasm Enhancement and Specialty Agriculture, Wuhan Botanical Garden, Chinese Academy of Sciences, Wuhan 430074, Hubei, China; Center of Conservation Biology, Core Botanical Gardens, Chinese Academy of Sciences, Wuhan 430074, Hubei, China; University of Chinese Academy of Sciences, Beijing, 100049 China; School of Integrative Plant Science, Section of Plant Biology and the L.H. Bailey Hortorium, Cornell University, Ithaca, NY 14850, USA; BTI Computational Biology Center, Boyce Thompson Institute, Ithaca, NY 14853, USA; Key Laboratory for Plant Diversity and Biogeography of East Asia, Kunming Institute of Botany, Chinese Academy of Sciences, Kunming, Yunnan, 650201 China

**Author notes:** Authors for correspondence: Tao Deng, Hengchang Wang, Hang Sun. These authors contributed equally to this work (Xu Zhang, Jacob B. Landis).

**Keywords:** radiating diversification, *Saussurea*, the Qinghai-Tibet Plateau, biodiversity hotspots, adaptive traits, diversification rates, ecological niche

## Abstract

- The Qinghai-Tibet Plateau (QTP) encompasses areas with a remarkably high degree of biodiversity, harboring exceptional species-rich radiations. How these radiations formed by interacting with geology, climate and ecology remains seldom examined.
- We investigate the roles of abiotic (environmental) and biotic (species-intrinsic) factors in driving radiating diversification of *Saussurea* (Asteraceae) by deploying a number of time-dependent, paleoenvironment-dependent and trait-dependent models, as well as ecological distribution data.
- We show that three main clades of *Saussurea* begin to diversify in the Miocene almost simultaneously, with increasing diversification rates toward the present and negative dependence to paleotemperature. Acceleration in diversification rates are correlated with adaptive traits, as well climate lability, niche breadth and species range.
- We conclude that fluctuation of paleoclimate along with complex QTP environments provided opportunities for increased diversification rates of *Saussurea* with diverse adaptive traits, highlighting the importance of combinations of clade-specific traits and ecological niches in driving rapid radiation.

## Introduction

The diversification pattern of species-rich rapid radiations reflects the evolutionary dynamics of biodiversity hotspots (Linder & Verboom, 2015). Understanding how these radiating lineages formed in response to historical process can advance our knowledge of adaptive evolution and enhance our ability to predict the threats to biodiversity posed by global warming (Ding *et al*., 2020). Mountainous regions represent just one-eighth of terrestrial land surface but are home to one-third of all species and exceptional species-rich radiations (Antonelli, 2015; Schwery *et al*., 2015; Antonelli *et al*., 2018). Particularly enigmatic is the Qinghai-Tibet Plateau (QTP) region, also known as the “Third Pole,” characterized by a complex geographical history and encompassing areas of remarkably high degree of biodiversity (Favre *et al*., 2015; Xing & Ree, 2017; Chen *et al*., 2018; Ding *et al*., 2020; Spicer *et al*., 2020). The QTP stands out as the earth’s highest and largest plateau, and includes the Himalaya and Hengduan Mountains which are listed as two of the 36 hotspots of biodiversity in the world (Myers *et al*., 2000; Li *et al*., 2014; Wen *et al*., 2014; Favre *et al*., 2015). The presence of steep environmental gradients in temperature and precipitation create abundant micro-habitats providing a variety of ecological niches essential for evolutionary radiations on the QTP (Mosbrugger *et al*., 2018; Muellner-Riehl *et al*., 2019). While a plethora of studies have suggested that diversification of plants on the QTP have evolved in association with plateau uplifting processes (reviewed by Wen *et al*., 2014), how such high species diversity form in such a short period of geologic time, and the interactions with geography, climate and ecology, remain seldom examined.

Evolutionary and diversification patterns of plants are often correlated with environmental abiotic forces, such as abrupt changes in climate or geological tectonic events that drive speciation and extinction rates, and/or species-intrinsic/biotic factors, such as interactions among species and key innovation traits (Drummond *et al*., 2012; Hughes & Atchison, 2015; Condamine *et al*., 2018; Muellner-Riehl *et al*., 2019; Nürk *et al*., 2019). There is a gap in our current understanding of radiating diversification drivers in the flora of the QTP, with previous studies mostly providing only a temporal (molecular dating) framework associating rapid radiations with the time span of plateau uplifting (e.g. Wang *et al*., 2009; Zhang *et al*., 2014; Xu *et al*., 2019). Employing models assuming continuous variation in diversification rates over time that depend on paleoenvironmental variables is essential to precisely determine how diversification rates are affected by abiotic environmental changes (Condamine *et al*., 2013; Sun *et al*., 2020). In addition to abiotic factors, diversification shifts are often correlated with the evolution of certain functional traits (Hughes & Atchison, 2015). Examples include geophytism in monocots leading to higher rates of diversification (Howard *et al*., 2020), polyploidization promoting species diversification of *Allium* (Han *et al*., 2020), and pollinator shifts, fruit types as well as elevational changes in the Andean bellflowers (Lagomarsino *et al*., 2016). Furthermore, the inclusion of ecological niche data is also crucial, because this reflects the interplay between historical processes and species intrinsic factors (Lavergne *et al*., 2010; Folk *et al*., 2019; Muellner-Riehl *et al*., 2019).

Here, we address the knowledge gap of rapid diversification by examining the roles of abiotic (environmental) and biotic (species-intrinsic) factors in driving radiating diversification of the species-rich genus *Saussurea* DC. (Asteraceae). *Saussurea* is one of the most diverse genera in Asteraceae, serving as an ideal study system for investigating the evolutionary patterns of a rapid radiation. The genus comprises approximately 400 species that are distributed in Asia, Europe and North America, with the highest diversity in the QTP (Wang *et al*., 2009; Shi & Raab-Straube, 2011; Chen, 2015; Zhang, *et al*., 2019a). Uncertainty in the number of species has largely been attributed to the complex taxonomy of related QTP taxa (Chen & Yuan, 2015), indicative of a recent radiation.

*Saussurea* exhibits extraordinary morphological diversity. For example, the most impressive species groups are the ‘snowball plants’ or ‘snow rabbits’, *S*. subg. *Eriocoryne*, with a thick woolly indumentum (densely haired), and the so-called ‘greenhouse plants’ or ‘snow lotuses’, *S*. subg. *Amphilaena*, in which the synflorescence is hidden by semi-transparent, white, yellowish or purple leafy bracts (Shi & Raab-Straube, 2011; Chen, 2015). *Saussurea* is present in virtually all possible habitats of the QTP, including steppes, moist forests, cold and dry alpine meadows, and scree slopes above 5,000 m, demonstrating a highly adaptive capability (Shi & Raab-Straube, 2011). Previous studies suggested that attractive morphological traits were the result of convergent adaptation to diverse environments in the QTP (Kita *et al*., 2004; Wang *et al*., 2009; Zhang, *et al*., 2019a), yet their contributions to the high-level diversity of *Saussurea* are still elusive. While biogeographic analysis inferred that *Saussurea* arose during the Miocene in the Hengduan Mountains (Xu *et al*., 2019), limited information about macro-evolutionary patterns related to historical climate and geologic processes were provided due to the lack of modeling diversification rates.

A robust phylogenetic framework is the basis for large-scale analyses of evolutionary patterns (Koenen *et al*., 2020), yet previous studies mainly relied on fragment DNA markers (e.g. Han *et al*., 2020; Howard *et al*., 2020; Sun *et al*., 2020), which have been revealed to provide insufficient phylogenetic signals and always yield parallel relationships for phylogenies of rapid radiations (Whitfield & Lockhart, 2007; Wang *et al*., 2009). In the present study, we reconstructed a robust time-calibrated phylogeny of *Saussurea* using 226 complete plastomes to explore the role played by abiotic and biotic factors in this rapidly radiating clade. If evolutionary dynamics are driven primarily by abrupt abiotic perturbations, we would expect diversification rate shifts following major climate changes that extirpated certain lineages while favoring the radiation of others. In contrast, if biotic factors or interactions among species are the dominant drivers of evolution, we would expect diversification shifts to be correlated with the evolution of functional traits and/or the colonization of new habitats (Condamine *et al*., 2018). While in a joint-effect scenario, diversification rates may vary continuously through time and paleoenvironments may shift with some clade-specific traits. We could hypothesize that fluctuations of terrestrial and climatic systems provide vast ecological opportunities, which are seized by lineages with ample adaptive traits and promote rapid radiating, emphasizing the decisive role of morphological diversity/plasticity and ecological niche availability. To test these hypotheses, we deployed a number of time-dependent, paleoenvironment-dependent and trait-dependent models, as well as ecological distribution data. Our study is designed to address the effects of paleoenvironmental and biological drivers on radiating diversification in the biodiversity hotspots, while providing a compelling example of the pivotal roles of morphological diversity and ecological niche.

## Materials and Methods

### Plastome Sampling, Sequencing and Assembly

To build a dated phylogeny of the genus *Saussurea*, we newly sequenced plastomes for 63 species and downloaded 163 additional plastomes from GenBank (accessed 29 November 2019); collectively these species included 199 taxa of *Saussurea* and 27 outgroup taxa. Collection details of the specimens were provided in Supporting Information Table S1. Total genomic DNA of was extracted from silica-gel dried leaves with a modified hexadecyltrimethylammonium bromide (CTAB) method (Yang *et al*., 2014). Purified DNA was fragmented and used to construct short-insert (500 bp) libraries per the manufacturer’s instructions (Illumina). Libraries were quantified using an Agilent 2100 Bioanalyzer (Agilent Technologies, Santa Clara, CA, USA), and were then sequenced on an Illumina HiSeq 4000 platform at Novogene Co., Ltd. in Kunming, Yunnan, China. Raw reads were directly assembled with the organellar assembler NOVOPLASTY v.2.7.2 (Dierckxsens *et al*., 2017), using a seed-and-extend algorithm employing the plastome sequence of *Saussurea japonica* (Genbank accession: MH926107.1) as the seed input, and all other parameters kept at default settings.

Assembled plastome sequences were initially annotated using Plastid Genome Annotator (PGA) (Qu *et al*., 2019), and then manually checked in GENEIOUS v.9.0.5 (Kearse *et al*., 2012).

### Estimates of Divergence Times

Our prior study suggested that including noncoding regions in phylogenetic analysis can maximize the power to resolve relationships of *Saussurea* (Zhang, *et al*., 2019a). Whole plastome sequences of 226 samples containing one inverted repeat region were aligned using MAFFT v.7.22 (Katoh & Standley, 2013). Poorly aligned regions were removed with TRIMAL v.1.2 (Capella-Gutiérrez *et al*., 2009) using the command ‘-automated1’. Age estimates were obtained using Markov Chain Monte Carlo (MCMC) analysis in BEAST v.1.10.4 (Suchard *et al*., 2018). We used a GTR + I + Γ nucleotide substitution model, uncorrelated relaxed lognormal clock and a birth-death model for the tree prior (Suchard *et al*., 2018). The MCMC analysis was run for 500 million generations, sampling every 10,000 generations, resulting in 50,000 samples in the posterior distribution of which the first 10,000 samples were discarded as burn-in. Convergence and performance of the MCMC runs were checked using TRACER v.1.6 (Rambaut *et al*., 2018). A maximum clade credibility (MCC) tree was then reconstructed in TREEANNOTATOR v.1.8.4 (Rambaut & Drummond, 2010), with median age and 95% height posterior density (HPD) annotated. Two high confident fossil calibrations with lognormal distributions were assigned: (A) The crown age of *Carduus-Cirsium* group was set to a minimum age of 14 million years ago (Mya) based on the Middle Miocene achenes identified as *Cirsium* (Mai, 1995; Barres *et al*., 2013); (B) The split of *Centaurea* and *Carthamus* was calibrated with a minimum age of 5 Mya, based on the records of pollen and achenes for *Centaurea* dating from the Early Pliocene (Popescu, 2002).

Additionally, the crown age of Cardueae was set to 39.2 Mya as a secondary calibration with a normal distribution based on the estimation by Barres *et al*. (2013).

### Estimates of Diversification rate

We explored the diversification dynamics of *Saussurea* using BAMM 2.5.0 (Rabosky, 2014), which employs a reversible-jump MCMC to sample a large number of possible diversification regimes from a given time-calibrated phylogeny. We pruned the MCC tree for BAMM analysis and retained only one sample of each species. Prior values were selected using the ‘setBAMMpriors’ function in the R package BAMMtools v.2.1.7 (R Core Team, 2014; Rabosky *et al*., 2014). Due to the controversial species number in *Saussurea*, the incomplete taxon sampling was appropriately set as 0.5 for all following analyses. The MCMC was run for 500 million generations and sampled every 50,000 generations. Post-run analyses were performed using the BAMMtools, with an initial 10% of the MCMC run discarded as burn-in, and the remaining data assessed for convergence and ESS values > 200. Rates-through-time plots were generated using ‘PlotRateThroughTime’ function for the entire genus as well as three clades. Speciation rates of *Saussurea* species were obtained using the ‘getTipRates’ function. Considering recent criticism relating to the statistical methods for lineage specific diversification models like BAMM (Moore *et al*., 2016; but also see Rabosky *et al*., 2017), we also employed the semiparametric DR statistic to calculate speciation rates, following the method described in Jetz *et al*. (2012) and Sun *et al*. (2020). Analysis of variance (ANOVA) was performed to determine whether differences among three phylogenetic clades and among four traditional subgenera were significant. In addition, we used TESS v.2.1 (Höhna *et al*., 2016) in R to detect the abrupt changes in speciation and extinction rates, applying the R-scripts of Condamine *et al*. (2018).

### Paleoenvironment dependent analyses

To quantify the effects of past environmental conditions on *Saussurea* diversification, we used RPANDA v1.9 (Condamine *et al*., 2013) to fit a series of time- and temperature-dependent likelihood diversification birth-death (BD) models, following the methodology of Condamine *et al*. (2018). Briefly, seven models were tested: BD model with constant λ (speciation rate) and μ (extinction rate) (i); BD model with λ dependent to time (ii) and environment (iii) exponentially, and constant μ; BD model with constant λ, and μ dependent to time (iv) and environment (v) exponentially; and BD model with λ and μ dependent to time (vi) and environment (vii) exponentially. Thus, we can obtain the equations: λ(E)= λ0 ×e^αE^ and μ(E)= μ0×e^βE^, in which λ0 and μ0 are the speciation and extinction rates for a given environmental variable. The values of α and β are the rates of change according to the environment, and positive values for them mean a positive effect of the environment on speciation or extinction (Condamine *et al*., 2013). We used paleotemperature over the last 12 million years (Myrs) (retrieved from Zachos *et al*., 2008) as environmental variables, and randomly sampled 500 trees from the BEAST posterior distribution (outgroups removed) to accommodate phylogenetic and dating uncertainties. The R package PSPLINE v.1.0 (Ben & Roberto, 2008) was used to visualize the speciation rates variating with paleoenvironmental variables.

### Trait dependent analyses

Nine characters were selected and coded based on descriptions in eFloras (http://www.efloras.org/), herbarium specimens and taxonomic literature, or were manually checked directly using online herbarium specimens from the Chinese Virtual Herbarium (http://www.cvh.ac.cn/), JSTOR (https://plants.jstor.org/), and field collections (Supporting Information Table S2). These characters included four binary morphological traits: stemless (0) vs. cauliferous (1), stem glabrous (0) vs. densely haired (1), the absence (0) vs. presence (1) of leafy bracts, and capitula solitary (0) vs. numerous (1); four multistate morphological traits: leaf margin entire (1) vs. pinnately lobed (2) vs. both types (3), leaves glabrous (1) vs. sparsely haired (2) vs. densely haired (3), phyllary in <5 (1) vs. 5 (2) vs. 6 (3) vs. >6 (4) rows, and phyllary glabrous (1) vs. sparsely haired (2) vs. densely haired (3) vs. appendage (4); as well as the geographical habitats: widespread (0) vs. alpine (1) vs. lowland (2).

The diversification rate shifts of binary traits were investigated using the hidden state speciation and extinction (HiSSE) model, which allows us to demonstrate hypotheses related to both the effects of the observed traits as well as incorporate unmeasured factors (Beaulieu & O’Meara, 2016). As described in Beaulieu and O’Meara (2016), 25 models were tested in the R package HISSE v.1.9.10: a full HiSSE model allowing all states to vary independently; four binary state speciation and extinction (BiSSE)-like models that excluded hidden states or constrained specific parameters of λ, μ, and transition rates (q); four null HiSSE models with various character-independent diversification (CID) forms; and 16 models assuming a hidden state associated with both observed character states with a variety of constrained values for λ, μ, and q (Supporting Information Table S3).

The best-fitting model was selected based on likelihood-ratio tests under a Chi-square distribution and Akaike’s information criterion (AIC) (Akaike, 1974). We also used a nonparametric FiSSE model (Fast, intuitive SSE model; Rabosky & Goldberg, 2017) serving as a complement to measure the robustness of our results. For multistate traits, MuSSE analyses were performed in the R package DIVERSITREE v.0.9.10 (FitzJohn, 2012) by fitting four distinct models with subsequent ANOVA testing: a null model with fully constrained variables; a full model allowing all variables to change independently; a model constraining each μ to be equal (free λ); and a model constraining the λ values for each state to be equal (free μ). Further estimates for the parameters of λ, μ, and net diversification rates (λ - μ) for each state were obtained in a Bayesian approach by incorporating a MCMC analysis with an exponential prior with 5,000 generations. A GeoHiSSE analysis was used to test hypotheses about range-dependent diversification processes (Caetano *et al*., 2018), implemented in the HISSE package. Two models with a range-independent diversification process and two other models in which the range have an effect on the diversification rate of the lineages were deployed and compared, as described in Caetano *et al*. (2018).

### Ecological distribution data and association with diversification rates

We used the ‘occ_search’ function of RGBIF v.1.3.0 (Chamberlain & Boettiger, 2017) to retrieve GPS coordinates for *Saussurea* species from GBIF on October 28, 2020. We extracted only data records that were georeferenced and excluded any coordinates with zero and/or integer latitude and longitude. We then removed geographic outliers defined as boxplot outliers of species occurrences in R. Range size of each species was estimated by applying a five kilometer buffer around each locality point using the ‘gBuffer’ function of RGEOS v.0.5-5 (https://CRAN.R-project.org/package=rgeos) following the descriptions of Testo *et al*. (2019). Range size data were log-transformed before analysis to overcome their skewed distribution (Testo *et al*., 2019). We extracted the values of 19 bioclimatic variables (from 1970 to 2000) from WorldClim (http://worldclim.org) using RASTER v.2.6-6 (https://CRAN.R-project.org/package=raster), and calculated a mean value for each species. Highly correlated variables were identified with a Pearson’s correlation coefficient > 0.75, and were removed. The remaining eight most predicative bioclimatic variables were: annual temperature (BIO1), mean diurnal range (BIO2), isothermality (BIO3), max temperature of warmest month (BIO5), annual precipitation (BIO12), precipitation seasonality (BIO15), precipitation of warmest quarter (BIO18) and precipitation of coldest quarter (BIO19). The main variation of bioclimatic variables representing climate lability was estimated by extracting the first two principal components (PC1 and PC2) from a PCA in R. To calculate the ecological niche breadth, we first estimated environmental niche models (ENM) in the R package ENMTOOLS v.1.0.2 (Warren *et al*., 2010), and then measured the spatial heterogeneity of the distribution of suitability scores using Levins’ B metrics (Levins, 1968) (‘raster.breadth’ function).

To demonstrate whether ecological factors drove rapid diversification of *Saussurea* species, multiple QuaSSE tests were performed under different models using DIVERSITREE. Five models with increasing complexity were constructed to fit the changes in speciation rates with climate lability (PCs of bioclimatic variables), niche breadth and species range size. Moreover, we used the *ES-sim* tests (Harvey & Rabosky, 2018) to crosscheck the correlation pattern revealed by QuaSSE. In addition to the default inverse equal splits statistic (Harvey & Rabosky, 2018), the DR statistic was also used as a reliable estimator to investigate correlation between speciation rate and continuous ecological factors using the R-scripts retrieved from Sun *et al*. (2020).

### Data availability

All newly sequenced plastomes were deposited in the National Center for Biotechnology Information (NCBI) database with accession numbers provided in Supporting Information Table S1. R scripts used in this study are available on GitHub (https://github.com/ZhangXu-CAS/Saussurea-diversification.git).

## Results

### Divergence time and diversification rate

Our phylogeny resolved a median stem age of ca. 11.79 Mya (95% HPD, 8.38–15.35 Mya) for *Saussurea*, with three clades beginning to diversify in parallel during the Miocene (ca. 9.05 Mya, ca. 8.37 Mya and ca. 8.92 Mya, respectively; Figs 1a, Supporting Information Figs S1, S2), suggesting a rapid radiation in this period. Our tree topology showed that four traditional morphology-based subgenera of *Saussurea* are paraphyletic, indicating adaptive traits have occurred multiple times. BAMM analysis revealed a scenario in which two shifts in net diversification rates occurred within *Saussurea* with high posterior probability (Figs 1a, Supporting Information Figs S3). Rates-through-time plots showed that while slightly offset in timing, diversification rates of the three clades accelerated during the Pliocene (Figs 1a, 1b), when the temperature dropped sharply. BAMM tip rates showed that clade-3 (0.981 events Myr^−1^ per lineage) had significantly higher mean speciation rate than clade-2 (0.560 events Myr^−1^ per lineage) and clade-1 (0.708 events Myr^−1^ per lineage) (*p* < 0.001, Supporting Information Tables S4, S5). Among four morphological-based subgenera, speciation rates of *S*. subg. *Amphilaena* (0.945 events Myr^−1^ per lineage) was highest (*p* < 0.001, Fig 1c, Supporting Information Tables S4, S5). While DR statistic revealed no significant difference among three main clades (*p* = 0.099), and *S*. subg. *Saussurea* (1.106 events Myr^−1^ per lineage) have the highest mean speciation rate (*p* = 0.022, Supporting Information Fig S4, Tables S4, S6). TESS analysis suggested that speciation and exaction shifts had higher posterior probability during the Pleistocene, consistent with the BAMM results (Supporting Information Fig S5).

**Fig. 1.**
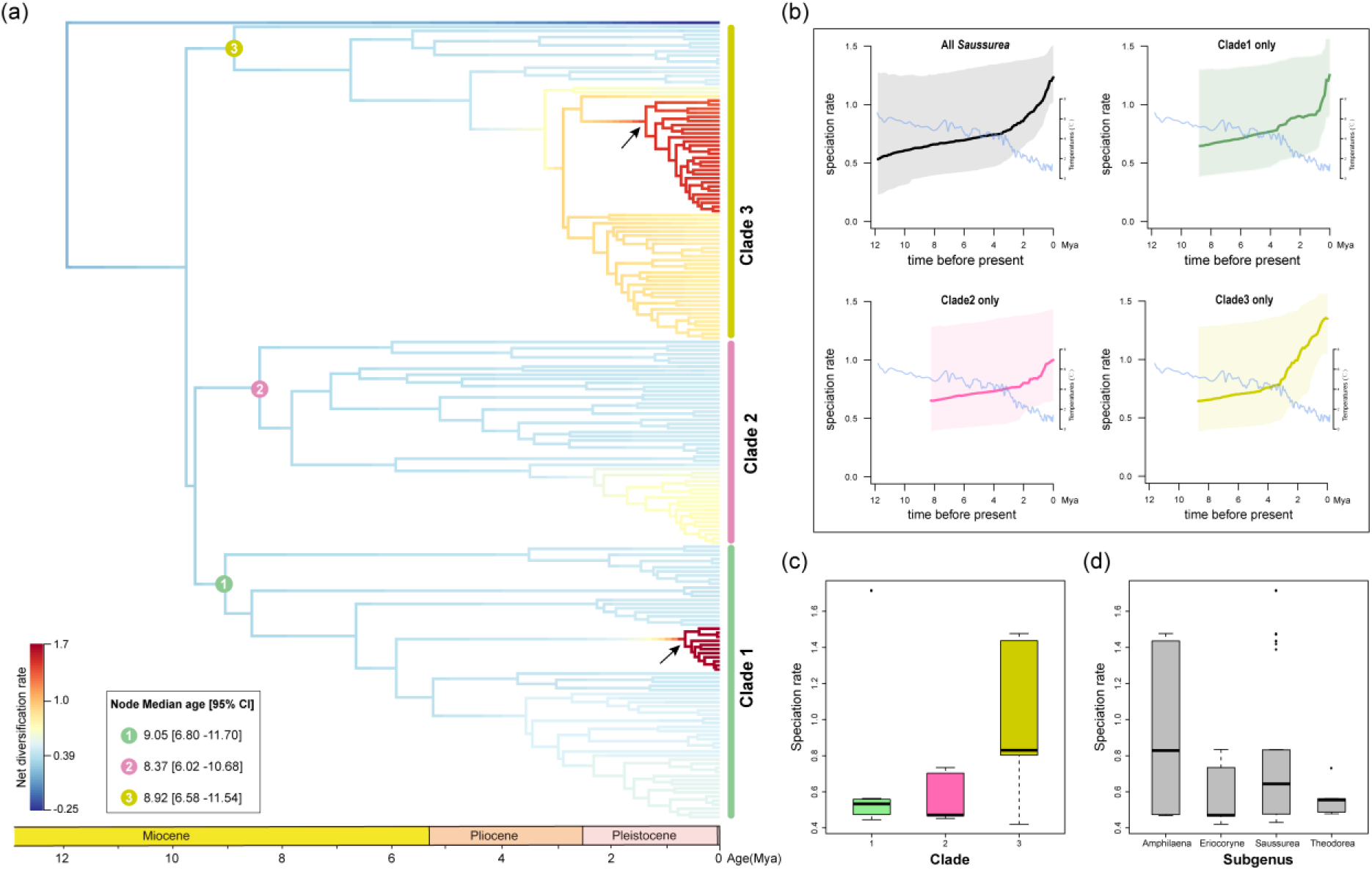
Diversification dynamics of *Saussurea* inferred from BAMM analysis. (a) BAMM identified two shifts in diversification rates (represented by arrows). The time of three clades beginning to diversify is provided. (b) Rates-through-time plots of all *Saussurea* species and three main clades separately, with trends in global climate change over 12 million years (Zachos *et al*. 2008) depicted. (c-d) BAMM tip rates of three clades and four morphology-based subgenera of *Saussurea*, respectively.

### Paleoenvironment dependent diversification

We used a maximum-likelihood framework to illustrate diversification dynamics dependent to paleoenvironment based on BD models to gain insight into the role of historical processes on diversification. Out of seven models, a model with temperature-dependent speciation fit the data best (Table 1). The best-fit model further indicated a negative dependence (α < 0) between past temperature and speciation rate for *Saussurea*, while extinction rate remained constant, suggesting extinction was likely not affected by temperature fluctuations. RPANDA results demonstrated a diversification regime in which diversification rates had opposite responses to changes of temperature over time, and accelerated sharply in the Pleistocene and increased toward the present (Fig. 2), consistent with the conclusion of rates-through-time in BAMM analysis (Fig. 1b).

**Table 1.** Results of RPANDA analyses.

| Models | NP | logL | AICc | $\lambda_0$ | $\alpha$ | $\mu_0$ | $\beta$ |
| --- | --- | --- | --- | --- | --- | --- | --- |
| Constant birth–death (1) | 2 | -325.7908 | 655.6424 | 0.7214 | NA | 0.3714 | NA |
| $\lambda_{\text{Time}}$ and $\mu_{\text{constant}}$ (2) | 3 | -325.2570 | 656.6364 | 0.6801 | -0.0618 | 0.1562 | NA |
| **$\lambda_{\text{Temp.}}$ and $\mu_{\text{constant}}$ (3)** | **3** | **-324.4240** | **654.9705** | **0.7585** | **-0.0933** | **0.1610** | **NA** |
| $\lambda_{\text{constant}}$ and $\mu_{\text{Time}}$ (4) | 3 | -325.4236 | 656.9698 | 0.7020 | NA | 0.3051 | 0.0475 |
| $\lambda_{\text{constant}}$ and $\mu_{\text{Temp.}}$ (5) | 3 | -325.4355 | 656.9930 | 0.649 | NA | 0.1445 | 0.2067 |
| $\lambda_{\text{Time}}$ and $\mu_{\text{Time}}$ (6) | 4 | -325.1840 | 658.5732 | 0.6840 | -0.0460 | 0.1843 | -0.0036 |
| $\lambda_{\text{Temp.}}$ and $\mu_{\text{Temp.}}$ (7) | 4 | -323.9815 | 656.168 | 0.693 | -0.0017 | 0.1910 | 0.1159 |

Bold columns represent the best model, in which speciation rate is negative dependence ( $\alpha < 0$ ) to past temperature and extinction rate is constant. Detailed model sets are described in Condamine *et al.* (2013). Abbreviations: NP, number of parameters; logL, log-likelihood; AICc, corrected Akaike Information Criterion. Parameter estimates: $\lambda_0$ and $\mu_0$ , speciation and extinction rates for a given environmental variable; and $\alpha$ , $\beta$ , parameter controlling variation of speciation and extinction with paleo-environment, respectively.

**Fig. 2.**
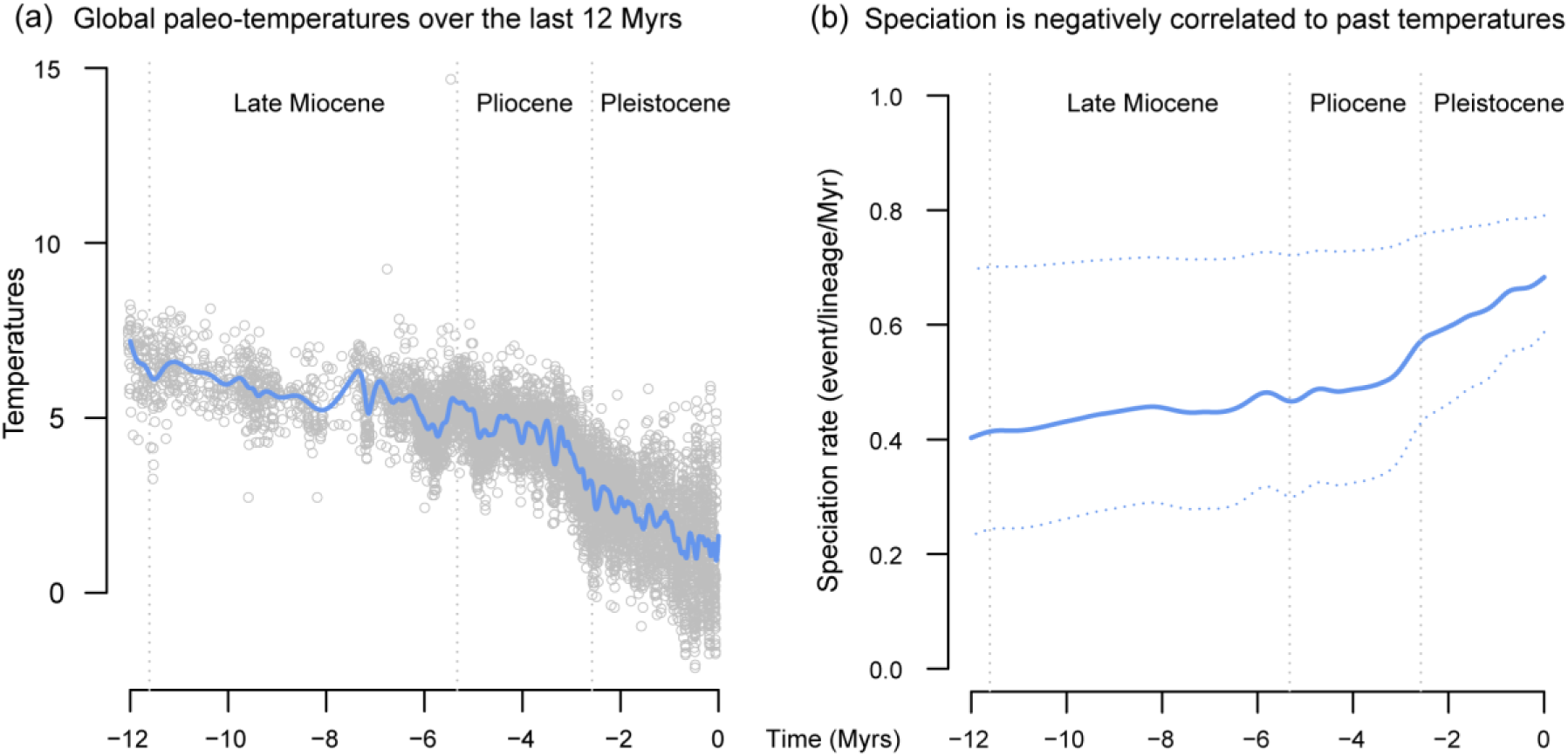
Paleoenvironment-dependent diversification processes in *Saussurea*. The best-fit paleoenvironment-dependent model implemented in RPANDA shows negative dependence between paleotemperatures (a) and speciation rate (b).

### Trait dependent diversification

We investigated eight morphological characters and geographical habitat that serve as a proxy for the effect of adaptive traits on diversification rate, to understand the role of trait innovations in the rapid radiation of *Saussurea*. For all four binary traits, the best model of the 25 models tested was the full HiSSE model with unique speciation, extinction and transition rates between the two character states observed and the hidden states (Supporting Information Tables S7). We then calculated mean speciation, extinction and net diversification rates values from the model-averaged marginal ancestral state reconstruction for each extant species in our tree. The results suggested that species with cauliferous plant, glabrous stem, leafy bracts and solitary capitula have higher mean speciation, extinction and net diversification (Table 2, Fig. 3). While the full HiSSE model showed observed differences in diversification rates between the states of these traits, it also indicated some unobserved traits drive the diversification. The complementary results from our FiSSE analysis supported the tendency of speciation rate revealed by HiSSE, but the only significant differences were between solitary capitula and numerous capitula (Table 2; *p* = 0.024).

**Table 2.** Summary of the mean rate values for four binary traits in HiSSE and FiSSE analysis.

| Trait | HiSSE |  |  | FiSSE |  |
| --- | --- | --- | --- | --- | --- |
| | $\lambda$ | $\mu$ | $\lambda-\mu$ | $\lambda$ | *p*-value |
| Stemless | 0.5947 | 0.2415 | 0.3532 | 0.9011 |  |
| **Cauliferous** | **1.0893** | **0.3821** | **0.4502** | **0.9264** | 0.4416 |
| **Glabrous** | **1.7230** | **1.2100** | **0.5128** | **0.8971** |  |
| Densely haired | 0.5531 | 0.2223 | 0.3308 | 0.8674 | 0.4096 |
| Normal | 0.8925 | 0.6849 | 0.2076 | 0.8987 |  |
| **Bracts** | **1.3951** | **0.8752** | **0.5199** | **0.9412** | 0.5614 |
| **Capitula solitary** | **1.6920** | **0.9969** | **0.6952** | **1.0825** |  |
| Capitula numerous | 0.5661 | 0.5253 | 0.0408 | 0.7828 | **0.0240** |

**Fig. 3.**
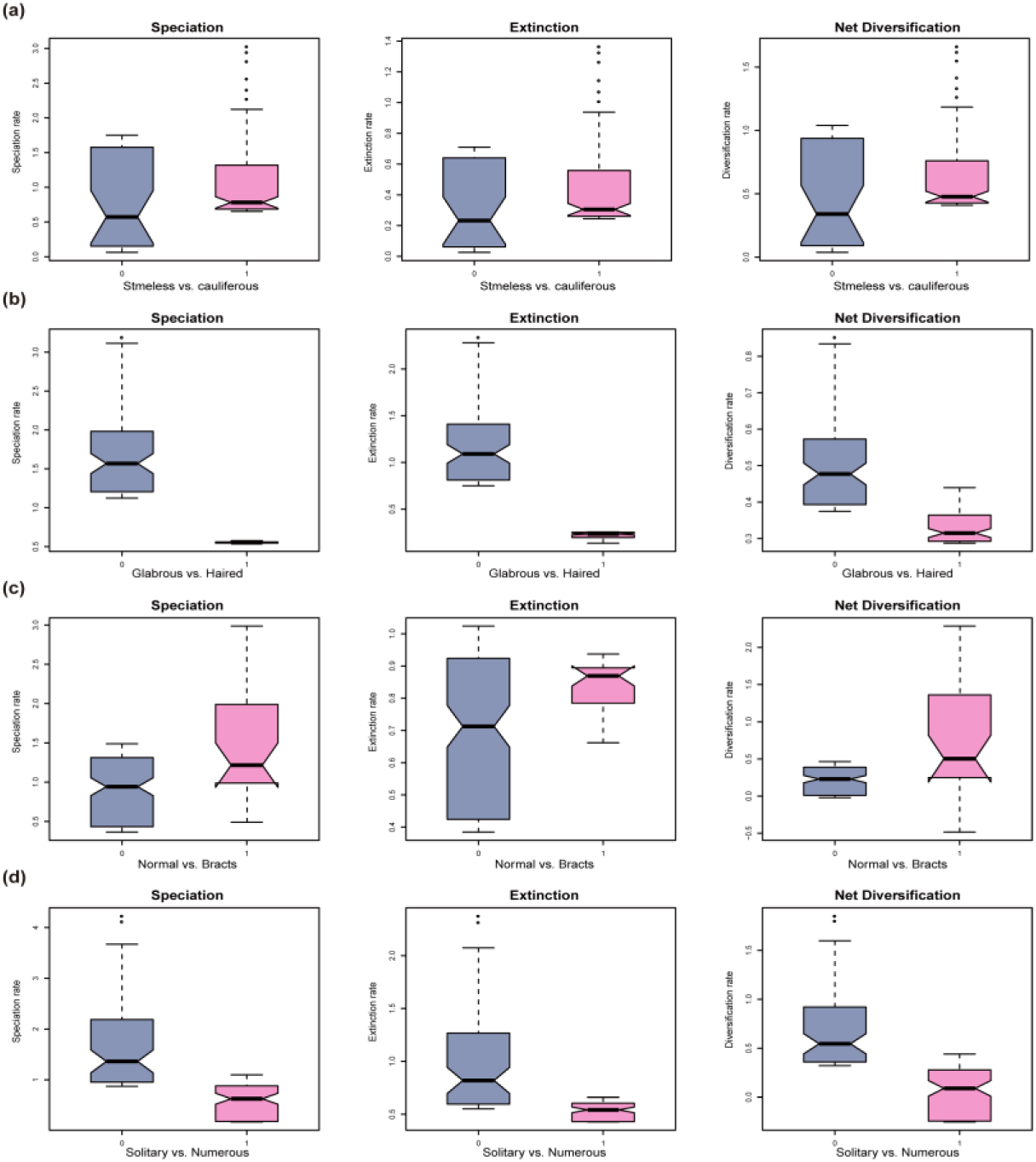
Binary trait dependent diversification of *Saussurea* inferred from HiSSE analysis. Speciation, extinction and net diversification rates are calculated by the model-averaged marginal ancestral state reconstruction for four binary traits: (a) stemless (0) vs. cauliferous (1), (b) stem glabrous (0) vs. densely haired (1), (c) the absence (0) vs. presence (1) of leafy bracts, and (d) capitula solitary (0) vs. numerous (1).

In the MuSSE analyses, ANOVA calculations all preferred models constraining each μ to be equal and allowing λ to vary (free λ), compared with either null models and full models (Supporting Information Table S8). The best-fitting model was then used as the starting point for a MCMC run of 5,000 generations to estimate the marginal distributions of rates for each traits using a state-dependent model (Fig. 4). Since all the models preferred constrained μ values, all of the estimated probability densities of μ overlapped (data not shown). The reconstructions of probability density of the net diversification rates (λ - μ) showed that some traits, i.e. leaf margin and phyllary types, have an overlap in net diversification rates among examined character states (Fig. 4a, d). The results suggest that species with glabrous leaves have higher net diversification rates than sparsely or densely haired species, consistent with higher mean rates for glabrous stem in the HiSSE analysis. For the phyllary character, the glabrous state also showed higher net diversification rates than sparsely or densely haired states despite some overlapping (Fig. 4d), and the rates of phyllary with six rows are higher than the remaining character states (Fig. 4c). The GeoHiSSE analysis suggested a model with one hidden area and no range-dependent diversification was the best fitting model (Supporting Information Table S9). From the result, we can see that species with lowland habitats have a substantially higher speciation, extinction and net diversification rates in comparison with both alpine and widespread distributions (Supporting Information Fig. S6).

**Fig. 4.**
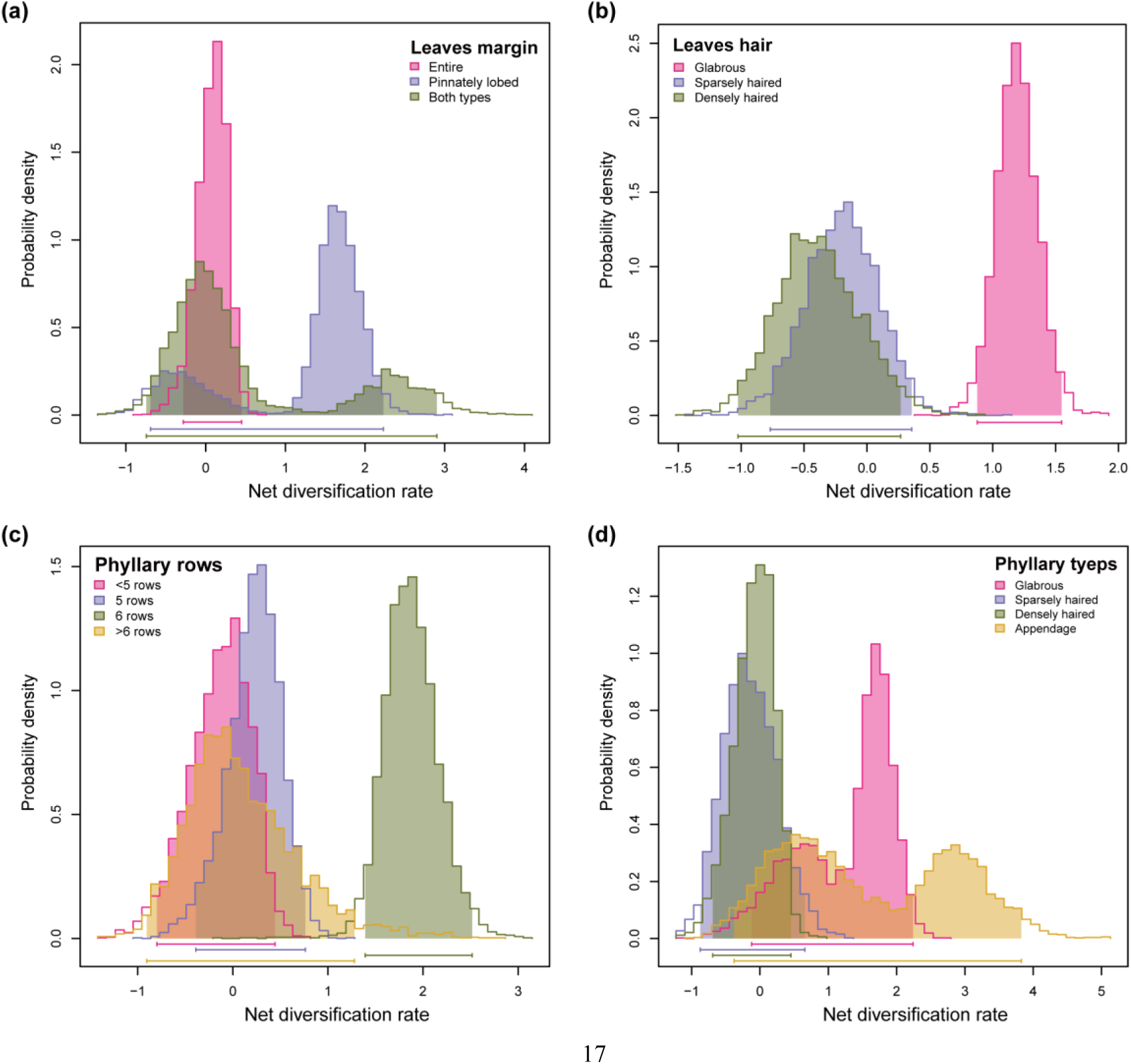
Multistate trait dependent diversification of *Saussurea* estimated from MuSSE analysis. Marginal distributions of net diversification rates are estimated by the MCMC run of 5, 000 generations for four multistate traits: (a) leaves margin entire (1) vs. pinnately lobed (2) vs. both types (3), (b) leaves glabrous (1) vs. sparsely haired (1) vs. densely haired (3), (c) phyllary in <5 (1) vs. 5 (2) vs. 6 (3) vs. >6 (4) rows, and (d) phyllary glabrous (1) vs. sparsely haired (2) vs. densely haired (3) vs. appendage (4).

### Ecological drivers of diversification

By correlating climate lability (PCs of bioclimatic variables), niche breadth and species range size with speciation rates (Supporting Information Table S10), we explored the role of ecological opportunities created by complex QTP environments in driving diversification of *Saussurea*. The first two PCs of bioclimatic variables explained 75.7% of the total climate variation in *Saussurea* (Supporting Information Fig. S7a). Among the eight retained bioclimatic variables, the precipitation of warmest quarter (BIO18) had the largest contribution to first two PCs, followed by the annual precipitation (BIO12) and the mean diurnal range (BIO2) (Supporting Information Fig. S7b). Under the QuaSSE analyses, PC1 of the climate variables showed a significant positive linear (l.m = 0.330, AIC = 1240.548, *p*-value = 0.005**) relationships with speciation rate, while climate PC2 preferred a constant model (AIC = 1183.524, *p*-value = 0.953); both niche breadth (AIC = 529.532, *p*-value < 0.000**) and species range size (AIC = 700.671, *p*-value < 0.000**) showed a significant positive sigmoidal (with drift) relationships with speciation rate (Supporting Information Table S11). Under the best sigmoidal models, the speciation rates of *Saussurea* kept a stable low state until the niche breadth and distribution range reached at 0.729 and 11.433 (log-transformed), respectively (midpoints; Fig. 5a, 5b). Under the *EM-sim* tests, both the DR statistic and the default inverse equal splits statistic revealed the same correlation pattern, in which niche breadth (ρ = 0.363 and 0.387, *p* = 0.027 and 0.019) and range size (ρ = 0.399 and 0.411, *p* = 0.018 and 0.011) showed significant positive relationship with speciation rates (Fig. 5c, 5d), while the correlation between speciation rates and climate lability (climate PC1: ρ = 0.170 and 0.188, *p* = 0.359 and 0.335; climate PC2: ρ = 0.098 and 0.095, *p* = 0.649 and 0.635) was not significant (Table 3).

**Table 3.**
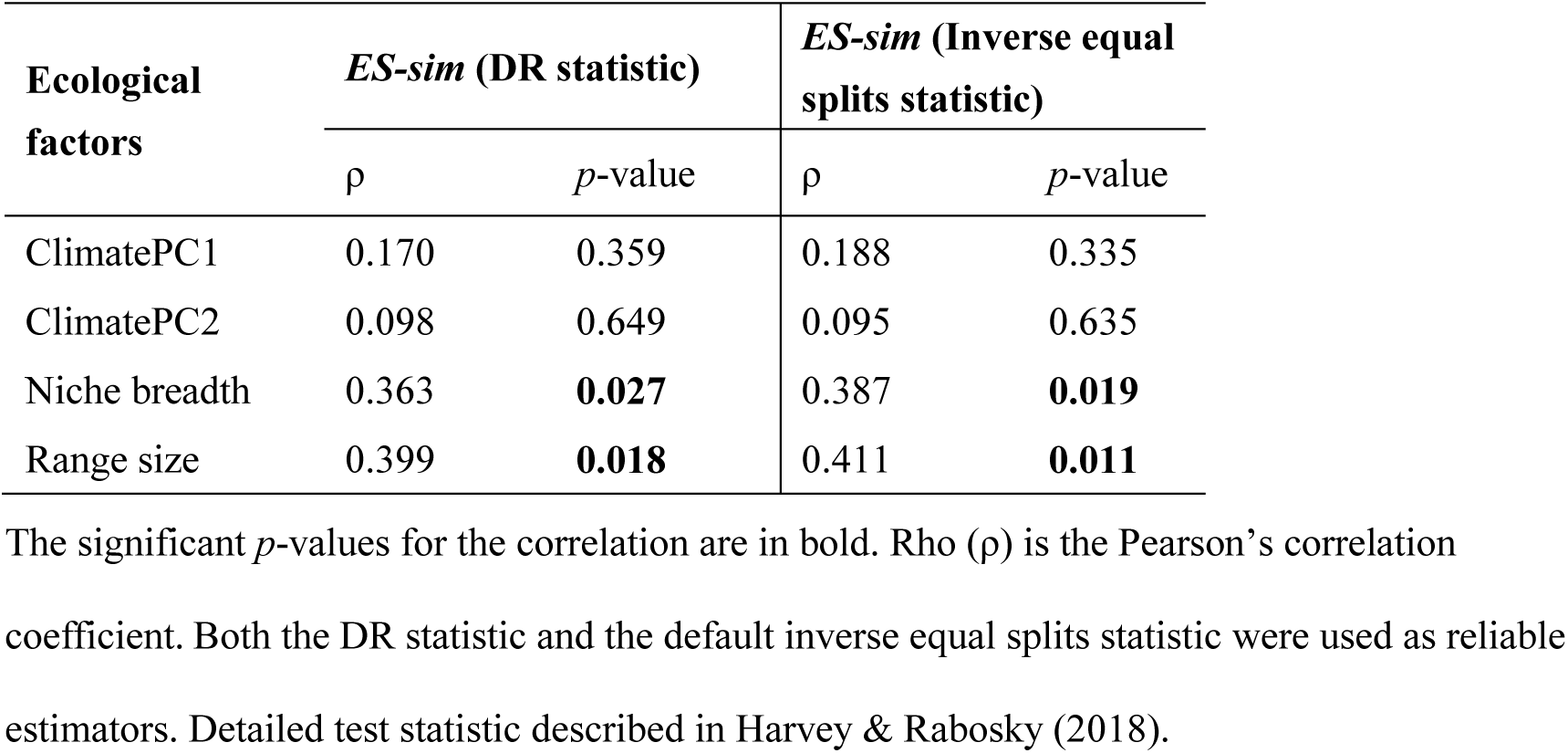
Summary of *ES-sim* tests for correlation between speciation rate and continuous ecological factors.

**Fig. 5.**
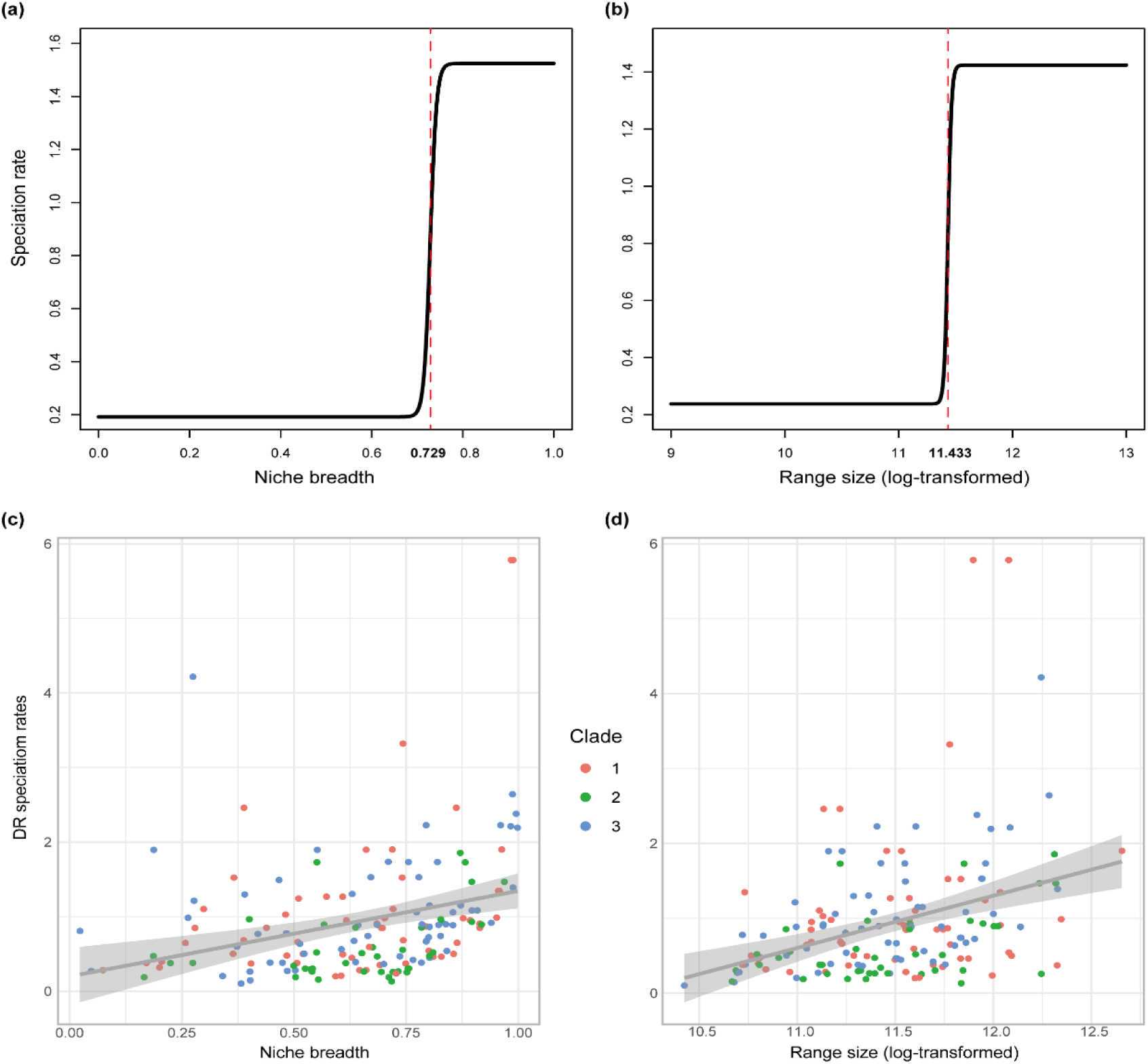
Speciation rates of *Saussurea* correlated with ecological factors based on the QuaSSE best-fitted model and *ES-sim* tests. Both (a) niche breadth and (b) species range size (log-transformed) show positive sigmoidal curves in QuaSSE analysis with the midpoints (represented by the red dashed line) of 0.729 and 11.433 on the x-axis respectively. *EM-sim* tests show significant positive relationships between DR speciation rates and (c) niche breadth and (d) species range size. Species from three clades are in different colors.

## Discussion

Our results demonstrate rapid diversification of *Saussurea* occurred in parallel during the Miocene, a period with extensive tectonic movement and climatic fluctuation on the QTP. A recent paper by (Louca & Pennell, 2020) raised limitations of macroevolutionary studies using estimated diversification rates, though several recent papers have suggested that more complex models (such as hidden state SSE models; (Helmstetter *et al*., 2021) and a hypothesis driven approach (Morlon *et al*., 2020) circumvent many of the issues raised. Therefore, we took an integrative approach to address the role that morphological traits and environmental conditions played in the evolutionary history of *Saussurea*. The rates of species diversification are revealed to be negatively correlated with paleotemperature, and accelerate sharply in the Pliocene toward the present. Similar patterns of increased diversification with global cooling have been documented in other flowering plant lineages, e.g. Saxifragales (Folk *et al*., 2019), rosids (Sun *et al*., 2020) and Campanulaceae (Lagomarsino *et al*., 2016), as well as in mammals (Stadler, 2011) and birds (Claramunt & Cracraft, 2015). Our trait dependent models detect some observed phenotypic adaptation associated with diversification changes, and indicate some unobserved traits also drive diversification, demonstrating a pivotal role of morphological diversity in this radiating diversification. Accounting for ecological niche data, we further reveal that acceleration in diversification rates are correlated with climate lability (PCs of bioclimatic variables), niche breadth and the size of species’ range. Overall, we conclude that tectonic activity of the QTP along with global paleoclimate cooling provided vast alpine niches for *Saussurea* species with ample adaptive traits, highlighting the important role of morphological diversity and ecological niche availability for species radiating to diverse environments.

We determined clade ages across *Saussurea* species using whole plastome sequences and found that the divergence of the main species clades occurred in the Miocene almost simultaneously. Compared to fragment DNA markers, plastomes have been shown to provide more sufficient phylogenetic signals which are powerful in resolving deep relationships of plant lineages (Parks *et al*., 2009; Wicke *et al*., 2011; Zhang *et al*., 2020). Our estimate for the origin of *Saussurea* (ca. 11.8 Ma) is consistent with the result from single-copy nuclear genes obtained via Hyb-Seq (ca. 12.5 Mya) (Herrando-Moraira *et al*., 2019) and the result from ITS sequences (12.6-10.3 Mya) (Wang *et al*., 2009), but was younger than the result of Xu *et al*. (2019) (ca. 18.5 Mya) using plastome coding regions and the result of Barres *et al*. (2013) (ca. 20.0 Mya) using chloroplast markers. The study of Barres *et al*. (2013) included only two species of *Saussurea* and used four chloroplast markers, *trnL-trnF, matK, ndhF* and *rbcL*. Different from Xu *et al*. (2019) setting the split of subtribe Arctiinae and subtribe Saussureinae as a minimum age to 8.0 Mya using the achene fossil assigned to *Arctium*, our study omitted this calibration because only one Arctiinae sample (*A. lappa*) was included in both studies and the relationship between Arctiinae and Saussureinae remains unresolved (Herrando-Moraira *et al*., 2019; Shen *et al*., 2020). In addition, we estimated divergence times using whole plastome sequences, as our prior work showed that including noncoding regions can maximize the resolution in resolving relationships of *Saussurea* (Zhang *et al*., 2019a).

Recent large-scale studies of species diversification on the QTP have provided convincing evidence for a Miocene diversification in plant lineages (Ding *et al*., 2020) as well as amphibians and reptiles (Xu *et al*., 2020). A hypothesis for the rich biodiversity found in mountainous regions like the QTP is uplift-driven diversification—that orogenic activities create diverse habitats favoring rapid *in situ* speciation of resident lineages (Xing & Ree, 2017; Chen *et al*., 2019). Extensive plateau uplift in the Miocene further intensified the Asian summer monsoon, which increased the precipitation for erosion through river incision, leading to greater topographic relief (Nie *et al*., 2018). This would have promoted the differentiation of microhabitats associated with elevational gradients and slope aspects, increasing the availability of ecological niches for radiating species (Ding *et al*., 2020). A previous study indicated that the *Saussurea* radiation was likely driven by ecological opportunities, similar to those on islands, provided by largely unoccupied habitats resulting from the extensive QTP uplifts (Wang *et al*., 2009). Our work provides compelling evidence of the vital role of ecological opportunities in *Saussurea* diversification by statistically correlating species niche breadth and distribution range to the speciation rate. A slight difference is that our result supports a wide-range radiation rather than an ‘island isolation’, from the positive correlation between range and speciation rate. We attribute the wide-range radiation of *Saussurea* to the presence of unique pappus combinations (Shi & Raab-Straube, 2011; Chen, 2015), which can promote the dispersal power of achenes to occupy more newly created niches. Therefore, colonizing success benefited by wide-range dispersal helped *Saussurea* species become one of the most diverse lineages on the QTP.

The negative correlation between paleotemperature and diversification rates in *Saussurea* does not seem surprising given the high species richness of *Saussurea* found at the high elevations of the QTP. Nonetheless, this insight is progressive for our understanding of the formation of the QTP flora, as it represents one of the few attempts to explicitly quantify the relationship between lineage diversification and a paleoenvironmental variable. Geological evidence suggests that after 15 Mya, global cooling and the further rise of QTP progressively led to more open, herb-rich vegetation as the modern high plateau formed with its cool, dry climate (Spicer *et al*., 2021). Thus, diversification among *Saussurea* clades could have been driven by increased ecological niches as suitable cold habitats became available. A sharply accelerated diversification rate of *Saussurea* was detected in the Pliocene toward the present. The uplift of the Hengduan Mountains region, at the southeastern margin of the QTP, is generally believed to have been rapid and recent, occurring mainly between the late Miocene and late Pliocene (Xing & Ree, 2017; Spicer *et al*., 2020). During the Quaternary glaciation, the Hengduan Mountains with its deep valleys would have provided numerous micro-refugia within the altitudinal and aspect heterogeneity (Sun *et al*., 2017; Spicer *et al*., 2021). This can explain why extensive morphological traits occur in parallel and evolved convergently, a result likely driven by local adaptation to the micro-habitats that were afforded by the complex and highly dissected landscape of the Hengduan Mountains.

Trait dependent analyses demonstrated that species exhibiting cauliferous plant, glabrous stem, leafy bracts and solitary capitula have higher speciation rates. These traits are usually observed in the subgenus *Amphilaena* (snow lotus), which is characterized by attractive leafy bract and is the symbols of snow mountains in the QTP (Shi & Raab-Straube, 2011; Chen, 2015). Snow lotus has abundant morphological variation and is a taxonomically complex group, with some new species described recently (e.g. Eckhard von, 2009; Chen & Yuan, 2015; Zhang, *et al*., 2019b). Despite having significant taxonomic characteristics, snow lotus is a non-monophyletic group, demonstrating that these adaptive traits have multiple origins and arose by convergent evolution. In fact, specialized leafy bracts, the so-called ‘glasshouse’ morphology, are prevalent among alpine species, such as in Lamiaceae, Asteraceae, and Polygonaceae (reviewed by Sun *et al*., 2014). Leafy bracts reportedly protect pollen grains from damage by UV-B radiation and rain, promote pollen germination by maintaining warmth, enhance pollinator visitation by providing a vivid visual display during flowering, and facilitate the development of fertilized ovules during seed development (Tsukaya, 2002; Yang & Sun, 2009; Song *et al*., 2015). Convergent morphological evolution seems to be common for plants adapting harsh environments of the QTP, examples include cushion (stemless) plant, woolly hairs and the leafy bract (Sun *et al*., 2014; Peng *et al*., 2015; Yang *et al*., 2019). Similar to leafy bract, the present of stemless and woolly hairs has been revealed to occurred multiple times, and is thought to defense cold and arid climate on the plateau (Sun *et al*., 2014). However, both stemless plants and the presence of woolly hairs appear to be not associated with an increase in diversification rate of *Saussurea*. A plausible explanation for this is that species with stemless and woolly hairs are commonly found in environments of the QTP with extremely high altitude with very low temperature, and these species usually have long lifespans.

Some traits associated with high diversification rates appear to have no evidence for ecological adaptation, such as solitary capitula and pinnate leaf margin. These may occur in combination with other important adaptive traits. Some traits were not examined because they are common across the entire genus, such as two rows of pappus and small achenes (Shi & Raab-Straube, 2011; Chen, 2015). Although trait dependent analyses showed several adaptive traits driving the increase of speciation rate, some unobserved traits were also important for rapid diversification, highlighting the vital roles of morphological diversity in the evolutionary history of *Saussurea*. Morphological diversity is an essential but often neglected aspect of biodiversity (Chartier *et al*., 2021). Our work provides a valuable guide for conservation efforts in the protection of morphological diversity of organisms, especially in the context of exacerbated biodiversity loss due to global warming.

Our results provided evidence of a positive relationship between speciation rate and niche breadth as well as species range. Among the few studies that have tested a niche breadth–diversification relationship, a clear consensus has not been reached (Sexton *et al*., 2017). One argument for low niche breadth lineages having greater diversification rates is that they are more likely to suffer from resource limitations and more susceptible to range fragmentation, and thus allopatric speciation occurs more frequently (Vrba, 1987). An alternative view is that species with high niche breadth typically have larger range sizes (Slatyer *et al*., 2013) and are therefore more likely to have these ranges fragmented by ecological or geographical barriers over evolutionary time, promoting allopatric speciation (Rolland & Salamin, 2016). We argue that wider ecological niches can help species diverging in the QTP cope with climatic fluctuation, occupy microhabitats and promote morphological divergence. Note that anthropogenic activities have led to landscape modification and habitat fragmentation, alternating the distributions of a vast array of species (Boivin *et al*., 2016), even in plateau areas (Chen *et al*., 2014). To promote future biodiversity resilience, the conservation of entire unfragmented landscapes is necessary to preserve niche heterogeneity and enable species migrations at will. Only this approach will conserve the processes of biodiversity dynamics as well as the genetic library and the capacity for future adaptation in threatened species (Spicer *et al*., 2020).

## Conclusion

Despite substantial processes on the taxonomy, phylogeny and biogeography of plant lineages on the QTP (reviewed by Wen *et al*., 2014), our knowledge of the diversification rates associated with geological activities along with subsequent environmental fluctuations and biotic interactions is still limited, especially for rapidly radiating species. Our study integrates *Saussurea* into an marcoevolutionary diversification framework. Using a genomic data set (plastome sequences) for reconstructing divergence history and multiple statistical analyses, we quantify the roles of abiotic/environmental and biotic/species-intrinsic factors in driving diversification of *Saussurea*. Our comprehensive and large-scale analyses depict a plausible evolutionary scene for *Saussurea*, and provide insights into the drivers of its radiating diversification. We document a Miocene diversification pattern in which increased speciation rates are related to global cooling, and correlate it to clade-specific traits and ecological niches. We hypothesize that the current mega diversity of *Saussurea* is the result of interactions between geological activity, global paleoclimate and ecological niche. Our results highlight the vital roles of morphological diversity and available ecological niches in plants adapting to the changing climate. Given the ongoing global warming and human expansion, causing the disappearance of numerous undescribed species and extensive occupied habitats, our present study together with previous macroevolutionary pattern studies (e.g. Condamine *et al*., 2018; Folk *et al*., 2019; Testo *et al*., 2019; Ding *et al*., 2020; Sun *et al*., 2020) provide valuable theoretical basis for mitigating the threats posed to biodiversity.

## Acknowledgements

We are grateful to all the collectors of *Saussurea* morphological data. We thank Ting-Shen Han for helpful in the visualization of Fig.5. We thank Jun-tong Chen and Kai Xue for providing the photos of *Saussurea* species. We also thank the members of the alpine research group of KIB, Jianwen Zhang, Zhuo Zhou, Hongliang Chen, Lishen Qian, Lu Sun and Yongzeng Zhang for helping with sample collection, and Yazhou Zhang for helping with species identification. This study was supported by the Second Tibetan Plateau Scientific Expedition and Research (STEP) program (2019QZKK0502), the Strategic Priority Research Program of Chinese Academy of Sciences (XDA20050203), the Key Projects of the Joint Fund of the National Natural Science Foundation of China (U1802232), the Major Program of the National Natural Science Foundation of China (31590823), the Youth Innovation Promotion Association of Chinese Academy of Sciences (2019382), the Young Academic and Technical Leader Raising Foundation of Yunnan Province (2019HB039), the Chinese Academy of Sciences “Light of West China“ Program, and the Biodiversity Survey, Monitoring and Assessment (2019HB2096001006), and the International Partnership Program of Chinese Academy of Sciences (151853KYSB20180009).

## Author contributions

HW, HS, TD and XZ developed the idea and designed the experiment; XZ and JBL performed the statistical analyses; XZ, JBL, TD, HS and HW interpreted the results and wrote the manuscript. XZ, YS, TF, HZ, NL, TB, XH and TD collected the leaf materials; All authors read, edited and approved the final manuscript.

## References

Akaike H. 1974. A new look at the statistical model identification. IEEE Transactions on Automatic Control 19: 716–723.

Antonelli A. 2015. Multiple origins of mountain life. Nature 524: 300–301.

Antonelli A, Kissling WD, Flantua SGA, Bermúdez MA, Mulch A, Muellner-Riehl AN, Kreft H, Linder HP, Badgley C, Fjeldså J, et al. 2018. Geological and climatic influences on mountain biodiversity. Nature Geoscience 11: 718–725.

Barres L, Sanmartin I, Anderson CL, Susanna A, Buerki S, Galbany-Casals M, Vilatersana R. 2013. Reconstructing the evolution and biogeographic history of tribe Cardueae (Compositae). American Journal of Botany 100: 867–882.

Beaulieu JM, O’Meara BC. 2016. Detecting hidden diversification shifts in models of trait-dependent speciation and extinction. Systematic Biology 65: 583–601.

Ben J, Roberto GG 2008. “PSPLINE: Stata module providing a penalized spline scatterplot smoother based on linear mixed model technology,” Statistical Software Components S456972, Boston College Department of Economics, revised 25 Jan 2009.

Boivin NL, Zeder MA, Fuller DQ, Crowther A, Larson G, Erlandson JM, Denham T, Petraglia MD. 2016. Ecological consequences of human niche construction: Examining long-term anthropogenic shaping of global species distributions. Proceedings of the National Academy of Sciences 113: 6388.

Caetano DS, O’Meara BC, Beaulieu JM. 2018. Hidden state models improve state-dependent diversification approaches, including biogeographical models. Evolution 72: 2308–2324.

Capella-Gutiérrez S, Silla-Martínez JM, Gabaldón T. 2009. trimAl: a tool for automated alignment trimming in large-scale phylogenetic analyses. Bioinformatics 25: 1972–1973.

Chamberlain SA, Boettiger C. 2017. R Python, and Ruby clients for GBIF species occurrence data. PeerJ Preprints 5: e3304v3301.

Chartier M, von Balthazar M, Sontag S, Löfstrand S, Palme T, Jabbour F, Sauquet H, Schönenberger J. 2021. Global patterns and a latitudinal gradient of flower disparity: perspectives from the angiosperm order Ericales. New Phytologist https://doi.org/10.1111/nph.17195.

Chen B, Zhang X, Tao J, Wu J, Wang J, Shi P, Zhang Y, Yu C. 2014. The impact of climate change and anthropogenic activities on alpine grassland over the Qinghai-Tibet Plateau. Agricultural and Forest Meteorology 189–190: 11-18.

Chen J, Huang Y, Brachi B, Yun Q, Zhang W, Lu W, Li H-n, Li W, Sun X, Wang G, et al. 2019. Genome-wide analysis of Cushion willow provides insights into alpine plant divergence in a biodiversity hotspot. Nature Communications 10: 5230.

Chen Y-S, Deng T, Zhou Z, Sun H. 2018. Is the East Asian flora ancient or not? National Science Review 5: 920–932.

Chen YS 2015. Asteraceae II *Saussurea*. In: Hong D-Y, Sun H, Watson M, Wen J, Zhang X-C eds. Flora of Pan-Himalaya. Beijing: Science Press.

Chen YS, Yuan Q. 2015. Twenty-six new species of *Saussurea* (Asteraceae, Cardueae) from the Qinghai-Tibetan Plateau and adjacent regions. Phytotaxa 213: 159–211.

Claramunt S, Cracraft J. 2015. A new time tree reveals Earth history’s imprint on the evolution of modern birds. Science Advances 1: e1501005.

Condamine FL, Rolland J, Höhna S, Sperling FAH, Sanmartín I. 2018. Testing the role of the red queen and court jester as drivers of the macroevolution of apollo butterflies. Systematic Biology 67: 940–964.

Condamine FL, Rolland J, Morlon H. 2013. Macroevolutionary perspectives to environmental change. Ecology Letters 16: 72–85.

Dierckxsens N, Mardulyn P, Smits G. 2017. NOVOPlasty: de novo assembly of organelle genomes from whole genome data. Nucleic Acids Res 45: e18.

Ding W-N, Ree RH, Spicer RA, Xing Y-W. 2020. Ancient orogenic and monsoon-driven assembly of the world’s richest temperate alpine flora. Science 369: 578.

Drummond CS, Eastwood RJ, Miotto STS, Hughes CE. 2012. Multiple Continental Radiations and Correlates of Diversification in *Lupinus* (Leguminosae): Testing for Key Innovation with Incomplete Taxon Sampling. Systematic Biology 61: 443–460.

Eckhard von R-S. 2009. *Saussurea luae* (Compositae, Cardueae), a new species of Snow Lotus from China. Willdenowia 39: 101–106.

Favre A, Päckert M, Pauls SU, Jähnig SC, Uhl D, Michalak I, Muellner-Riehl AN. 2015. The role of the uplift of the Qinghai-Tibetan Plateau for the evolution of Tibetan biotas. Biological Reviews 90: 236–253.

FitzJohn RG. 2012. DIVERSITREE: comparative phylogenetic analyses of diversification in R. Methods in Ecology and Evolution 3: 1084–1092.

Folk RA, Stubbs RL, Mort ME, Cellinese N, Allen JM, Soltis PS, Soltis DE, Guralnick RP. 2019. Rates of niche and phenotype evolution lag behind diversification in a temperate radiation. Proceedings of the National Academy of Sciences 116: 10874.

Han T-S, Zheng Q-J, Onstein RE, Rojas-Andrés BM, Hauenschild F, Muellner-Riehl AN, Xing Y-W. 2020. Polyploidy promotes species diversification of *Allium* through ecological shifts. New Phytologist 225: 571–583.

Harvey MG, Rabosky DL. 2018. Continuous traits and speciation rates: Alternatives to state-dependent diversification models. Methods in Ecology and Evolution 9: 984–993.

Helmstetter AJ, Glemin S, Käfer J, Zenil-Ferguson R, Sauquet H, de Boer H, Dagallier L- PMJ, Mazet N, Reboud EL, Couvreur TLP, et al. 2021. Pulled Diversification Rates, Lineages-Through-Time Plots and Modern Macroevolutionary Modelling. bioRxiv: 2021.2001.2004.424672.

Herrando-Moraira S, Calleja JA, Galbany-Casals M, Garcia-Jacas N, Liu JQ, López-Alvarado J, López-Pujol J, Mandel JR, Massó S, Montes-Moreno N, et al. 2019. Nuclear and plastid DNA phylogeny of tribe Cardueae (Compositae) with Hyb-Seq data: A new subtribal classification and a temporal diversification framework. Molecular Phylogenetics and Evolution 137: 313–332.

Höhna S, May MR, Moore BR. 2016. TESS: an R package for efficiently simulating phylogenetic trees and performing Bayesian inference of lineage diversification rates. Bioinformatics 32: 789–791.

Howard CC, Landis JB, Beaulieu JM, Cellinese N. 2020. Geophytism in monocots leads to higher rates of diversification. New Phytologist 225: 1023–1032.

Hughes CE, Atchison GW. 2015. The ubiquity of alpine plant radiations: from the Andes to the Hengduan Mountains. New Phytologist 207: 275–282.

Jetz W, Thomas GH, Joy JB, Hartmann K, Mooers AO. 2012. The global diversity of birds in space and time. Nature 491: 444–448.

Katoh K, Standley DM. 2013. MAFFT multiple sequence alignment software version 7: improvements in performance and usability. Molecular Biology and Evolution 30: 772–780.

Kearse M, Moir R, Wilson A, Stones-Havas S, Cheung M, Sturrock S, Buxton S, Cooper A, Markowitz S, Duran C, et al. 2012. Geneious Basic: an integrated and extendable desktop software platform for the organization and analysis of sequence data. Bioinformatics 28: 1647–1649.

Kita Y, Fujikawa K, Ito M, Ohba H, Kato M. 2004. Molecular phylogenetic analyses and systematics of the genus *Saussurea* and related genera (Asteraceae, Cardueae). Taxon 53: 679–690.

Koenen EJM, Ojeda DI, Steeves R, Migliore J, Bakker FT, Wieringa JJ, Kidner C, Hardy OJ, Pennington RT, Bruneau A, et al. 2020. Large-scale genomic sequence data resolve the deepest divergences in the legume phylogeny and support a near-simultaneous evolutionary origin of all six subfamilies. New Phytologist 225: 1355–1369.

Lagomarsino LP, Condamine FL, Antonelli A, Mulch A, Davis CC. 2016. The abiotic and biotic drivers of rapid diversification in Andean bellflowers (Campanulaceae). New Phytologist 210: 1430–1442.

Lavergne S, Mouquet N, Thuiller W, Ronce O. 2010. Biodiversity and Climate Change: Integrating Evolutionary and Ecological Responses of Species and Communities. Annual Review of Ecology, Evolution, and Systematics 41: 321–350.

Levins R. 1968. Evolution in changing environments. Princeton, NJ, USA: Princeton University Press.

Li X-H, Zhu X-X, Niu Y, Sun H. 2014. Phylogenetic clustering and overdispersion for alpine plants along elevational gradient in the Hengduan Mountains Region, southwest China. Journal of Systematics and Evolution 52: 280–288.

Linder HP, Verboom GA. 2015. The Evolution of Regional Species Richness: The History of the Southern African Flora. Annual Review of Ecology, Evolution, and Systematics 46: 393–412.

Louca S, Pennell MW. 2020. Extant timetrees are consistent with a myriad of diversification histories. Nature 580: 502–505.

Mai DH. 1995. Tertiäre Vegetationsgeschichte Europas : Methoden und Ergebnisse. Germany: Gustav Fischer Verlag.

Moore BR, Höhna S, May MR, Rannala B, Huelsenbeck JP. 2016. Critically evaluating the theory and performance of Bayesian analysis of macroevolutionary mixtures. Proceedings of the National Academy of Sciences 113.34: 9569–9574.

Morlon H, Hartig F, Robin S. 2020. Prior hypotheses or regularization allow inference of diversification histories from extant timetrees. bioRxiv: 2020.2007.2003.185074.

Mosbrugger V, Favre A, Muellner-Riehl AN, Päckert M, Mulch A 2018. Cenozoic Evolution of Geobiodiversity in the Tibeto-Himalayan Region. In: Hoorn C, Perrigo A, Antonelli A eds. Mountains, Climate and Biodiversity. UK: Wiley-Blackwell, 429–448.

Muellner-Riehl AN, Schnitzler J, Kissling WD, Mosbrugger V, Rijsdijk KF, Seijmonsbergen AC, Versteegh H, Favre A. 2019. Origins of global mountain plant biodiversity: Testing the ‘mountain-geobiodiversity hypothesis’. Journal of Biogeography 46: 2826–2838.

Myers N, Mittermeier RA, Mittermeier CG, da Fonseca GAB, Kent J. 2000. Biodiversity hotspots for conservation priorities. Nature 403: 853–858.

Nie J, Ruetenik G, Gallagher K, Hoke G, Garzione CN, Wang W, Stockli D, Hu X, Wang Z, Wang Y, et al. 2018. Rapid incision of the Mekong River in the middle Miocene linked to monsoonal precipitation. Nature Geoscience 11: 944–948.

Nürk NM, Atchison GW, Hughes CE. 2019. Island woodiness underpins accelerated disparification in plant radiations. New Phytologist 224: 518–531.

Parks M, Cronn R, Liston A. 2009. Increasing phylogenetic resolution at low taxonomic levels using massively parallel sequencing of chloroplast genomes. BMC Biology 7: 84.

Peng D-L, Niu Y, Song B, Chen J-G, Li Z-M, Yang Y, Sun H. 2015. Woolly and overlapping leaves dampen temperature fluctuations in reproductive organ of an alpine Himalayan forb. Journal of Plant Ecology 8: 159–165.

Popescu SM. 2002. Repetitive changes in Early Pliocene vegetation revealed by high-resolution pollen analysis: revised cyclostratigraphy of southwestern Romania. Review of Palaeobotany and Palynology 120: 181–202.

Qu XJ, Moore MJ, Li DZ, Yi TS. 2019. PGA: a software package for rapid, accurate, and flexible batch annotation of plastomes. Plant Methods 15: 50.

R Core Team 2014. R: A language and environment for statistical computing. R Foundation for Statistical Computing. Vienna, Austria.

Rabosky DL. 2014. Automatic Detection of Key Innovations, Rate Shifts, and Diversity-Dependence on Phylogenetic Trees. PLOS ONE 9: e89543.

Rabosky DL, Goldberg EE. 2017. FiSSE: A simple nonparametric test for the effects of a binary character on lineage diversification rates. Evolution 71: 1432–1442.

Rabosky DL, Grundler M, Anderson C, Title P, Shi JJ, Brown JW, Huang H, Larson JG. 2014. BAMMtools: an R package for the analysis of evolutionary dynamics on phylogenetic trees. Methods in Ecology and Evolution 5: 701–707.

Rabosky DL, Mitchell JS, Chang J. 2017. Is BAMM Flawed? Theoretical and Practical Concerns in the Analysis of Multi-Rate Diversification Models. Systematic Biology 66: 477–498.

Rambaut A, Drummond A. 2010. TreeAnnotator version 1.6. 1. University of Edinburgh, Edinburgh, UK: Available at: http://beast.bio.ed.ac.uk. [accessed 1 September 2019]

Rambaut A, Drummond AJ, Xie D, Baele G, Suchard MA. 2018. Posterior Summarization in Bayesian Phylogenetics Using Tracer 1.7. Systematic Biology 67: 901–904.

Rolland J, Salamin N. 2016. Niche width impacts vertebrate diversification. Global Ecology and Biogeography 25: 1252–1263.

Schwery O, Onstein RE, Bouchenak-Khelladi Y, Xing Y, Carter RJ, Linder HP. 2015. As old as the mountains: the radiations of the Ericaceae. New Phytologist 207: 355–367.

Sexton JP, Montiel J, Shay JE, Stephens MR, Slatyer RA. 2017. Evolution of Ecological Niche Breadth. Annual Review of Ecology, Evolution, and Systematics 48: 183–206.

Shen J, Zhang X, Landis JB, Zhang H, Deng T, Sun H, Wang H. 2020. Plastome Evolution in *Dolomiaea* (Asteraceae, Cardueae) Using Phylogenomic and Comparative Analyses. Frontiers in Plant Science 11: 376.

Shi Z, Raab-Straube Ev 2011. Cardueae. In: Wu ZY, Raven, P. H. & Hong, D. Y. ed. Flora of China. Beijing & St. Louis: Science Press & Missouri Botanical Garden Press, 42–194.

Slatyer RA, Hirst M, Sexton JP. 2013. Niche breadth predicts geographical range size: a general ecological pattern. Ecology Letters 16: 1104–1114.

Song B, Stöcklin J, Peng D, Gao Y, Sun H. 2015. The bracts of the alpine ‘glasshouse’ plant Rheum alexandrae (Polygonaceae) enhance reproductive fitness of its pollinating seed-consuming mutualist. Botanical Journal of the Linnean Society 179: 349–359.

Spicer RA, Farnsworth A, Su T. 2020. Cenozoic topography, monsoons and biodiversity conservation within the Tibetan Region: An evolving story. Plant Diversity 42: 229–254.

Spicer RA, Su T, Valdes PJ, Farnsworth A, Wu F-X, Shi G, Spicer TEV, Zhou Z. 2021. Why ‘the uplift of the Tibetan Plateau’ is a myth? National Science Review 8: nwaa091.

Stadler T. 2011. Mammalian phylogeny reveals recent diversification rate shifts. Proceedings of the National Academy of Sciences 108: 6187.

Suchard MA, Lemey P, Baele G, Ayres DL, Drummond AJ, Rambaut A. 2018. Bayesian phylogenetic and phylodynamic data integration using BEAST 1.10. Virus Evolution 4: vey016.

Sun H, Niu Y, Chen Y-S, Song B, Liu C-Q, Peng D-L, Chen J-G, Yang Y. 2014. Survival and reproduction of plant species in the Qinghai–Tibet Plateau. Journal of Systematics and Evolution 52: 378–396.

Sun H, Zhang J, Deng T, Boufford DE. 2017. Origins and evolution of plant diversity in the Hengduan Mountains, China. Plant Diversity 39: 161–166.

Sun M, Folk RA, Gitzendanner MA, Soltis PS, Chen Z, Soltis DE, Guralnick RP. 2020. Recent accelerated diversification in rosids occurred outside the tropics. Nature Communications 11: 3333.

Testo WL, Sessa E, Barrington DS. 2019. The rise of the Andes promoted rapid diversification in Neotropical *Phlegmariurus* (Lycopodiaceae). New Phytologist 222: 604–613.

Tsukaya H. 2002. Optical and anatomical characteristics of bracts from the Chinese “glasshouse” plant, *Rheum alexandrae* Batalin (Polygonaceae), in Yunnan, China. Journal of Plant Research 115: 59–63.

Vrba ES. 1987. Ecology in relation to speciation rates: some case histories of Miocene-Recent mammal clades. Evolutionary Ecology 1: 283–300.

Wang YJ, Susanna A, Von Raab-Straube E, Milne R, Liu JQ. 2009. Island-like radiation of *Saussurea* (Asteraceae: Cardueae) triggered by uplifts of the Qinghai-Tibetan Plateau. Biological Journal of the Linnean Society 97: 893–903.

Warren DL, Glor RE, Turelli M. 2010. ENMTools: a toolbox for comparative studies of environmental niche models. Ecography 33: 607–611.

Wen J, Zhang J, Nie Z-L, Zhong Y, Sun H. 2014. Evolutionary diversifications of plants on the Qinghai-Tibetan Plateau. Frontiers in Genetics 5: 4.

Whitfield JB, Lockhart PJ. 2007. Deciphering ancient rapid radiations. Trends in Ecology & Evolution 22: 258–265.

Wicke S, Schneeweiss GM, dePamphilis CW, Muller KF, Quandt D. 2011. The evolution of the plastid chromosome in land plants: gene content, gene order, gene function. Plant Molecular Biology 76: 273–297.

Xing Y, Ree RH. 2017. Uplift-driven diversification in the Hengduan Mountains, a temperate biodiversity hotspot. Proceedings of the National Academy of Sciences: 201616063.

Xu L-S, Herrando-Moraira S, Susanna A, Galbany-Casals M, Chen Y-S. 2019. Phylogeny, origin and dispersal of *Saussurea* (Asteraceae) based on chloroplast genome data. Molecular Phylogenetics and Evolution 141: 106613.

Xu W, Dong W-J, Fu T-T, Gao W, Lu C-Q, Yan F, Wu Y-H, Jiang K, Jin J-Q, Chen H-M, et al. 2020. Herpetological phylogeographic analyses support a Miocene focal point of Himalayan uplift and biological diversification. National Science Review nwaa263.

Yang J-B, Li D-Z, Li H-T. 2014. Highly effective sequencing whole chloroplast genomes of angiosperms by nine novel universal primer pairs. Molecular Ecology Resources 14: 1024–1031.

Yang Y, Chen J, Song B, Niu Y, Peng D, Zhang J, Deng T, Luo D, Ma X, Zhou Z. 2019. Advances in the studies of plant diversity and ecological adaptation in the subnival ecosystem of the Qinghai-Tibet Plateau. Chinese Science Bulletin 64: 2856–2864.

Yang Y, Sun H. 2009. The Bracts of *Saussurea velutina* (Asteraceae) Protect Inflorescences from Fluctuating Weather at High Elevations of the Hengduan Mountains, Southwestern China. Arctic, Antarctic, and Alpine Research 41: 515–521.

Zachos JC, Dickens GR, Zeebe RE. 2008. An early Cenozoic perspective on greenhouse warming and carbon-cycle dynamics. Nature 451: 279–283.

Zhang J-Q, Meng S-Y, Allen GA, Wen J, Rao G-Y. 2014. Rapid radiation and dispersal out of the Qinghai-Tibetan Plateau of an alpine plant lineage *Rhodiola* (Crassulaceae). Molecular Phylogenetics and Evolution 77: 147–158.

Zhang X, Deng T, Moore MJ, Ji Y, Lin N, Zhang H, Meng A, Wang H, Sun Y, Sun H. 2019a. Plastome phylogenomics of *Saussurea* (Asteraceae: Cardueae). BMC Plant Biology 19: 290.

Zhang X, Sun Y, Landis JB, Lv Z, Shen J, Zhang H, Lin N, Li L, Sun J, Deng T, et al. 2020. Plastome phylogenomic study of Gentianeae (Gentianaceae): widespread gene tree discordance and its association with evolutionary rate heterogeneity of plastid genes. BMC Plant Biology 20: 340.

Zhang Y, Tang R, Huang X, Sun W, Ma X, Sun H. 2019b. *Saussurea balangshanensis* sp. nov. (Asteraceae), from the Hengduan Mountains region, SW China. Nordic Journal of Botany 37: https://doi.org/10.1111/njb.02078.

