## Supporting Information Figures S1-S7, Tables S1-S911 for "Insights into the drivers of radiating diversification in biodiversity hotspots using *Saussurea* (Asteraceae) as a case"

**Fig. S1** Dated phylogeny of *Saussurea*.

**Fig. S2** Dated phylogeny showing the diversification time of three clades of *Saussurea*.

**Fig. S3** Summary of posterior probability of each shift rate in the BAMM analysis.

**Fig. S4** Speciation rates estimated by DR statistic.

**Fig. S5** Rate-through-time plot from the TESS analysis.

**Fig. S6** Range-dependent diversification of *Saussurea* inferred from GeoHiSSE analysis.

**Fig. S7** PCA results of eight bioclimatic variables representing climate lability of *Saussurea* species.

**Table S1** Sample information in the present study.

**Table S2** Traits coding regime in the present study.

**Table S3** HiSSE models used in the present study.

**Table S4** Speciation rates of Saussurea species estimated from BAMM and DR statistic.

**Table S5** Statistic summary of *Saussurea* speciation rates from BAMM analysis.

**Table S6** Statistic summary of *Saussurea* speciation rates from DR statistic.

**Table S7** Model comparison of HiSSE analysis for four binary traits of *Saussurea*.

**Table S8** Model comparison of MuSSE analysis for multistate traits of *Saussurea*.

**Table S9** Model comparison of GeoHiSSE analysis for ecological habitants of Saussurea.

**Table S10** Summary table of ecological factors for correlation with diversification rates of *Saussurea* species.

**Table S11** Model comparison of QuaSSE analysis for correlation between ecological factors and speciation rates of Saussurea

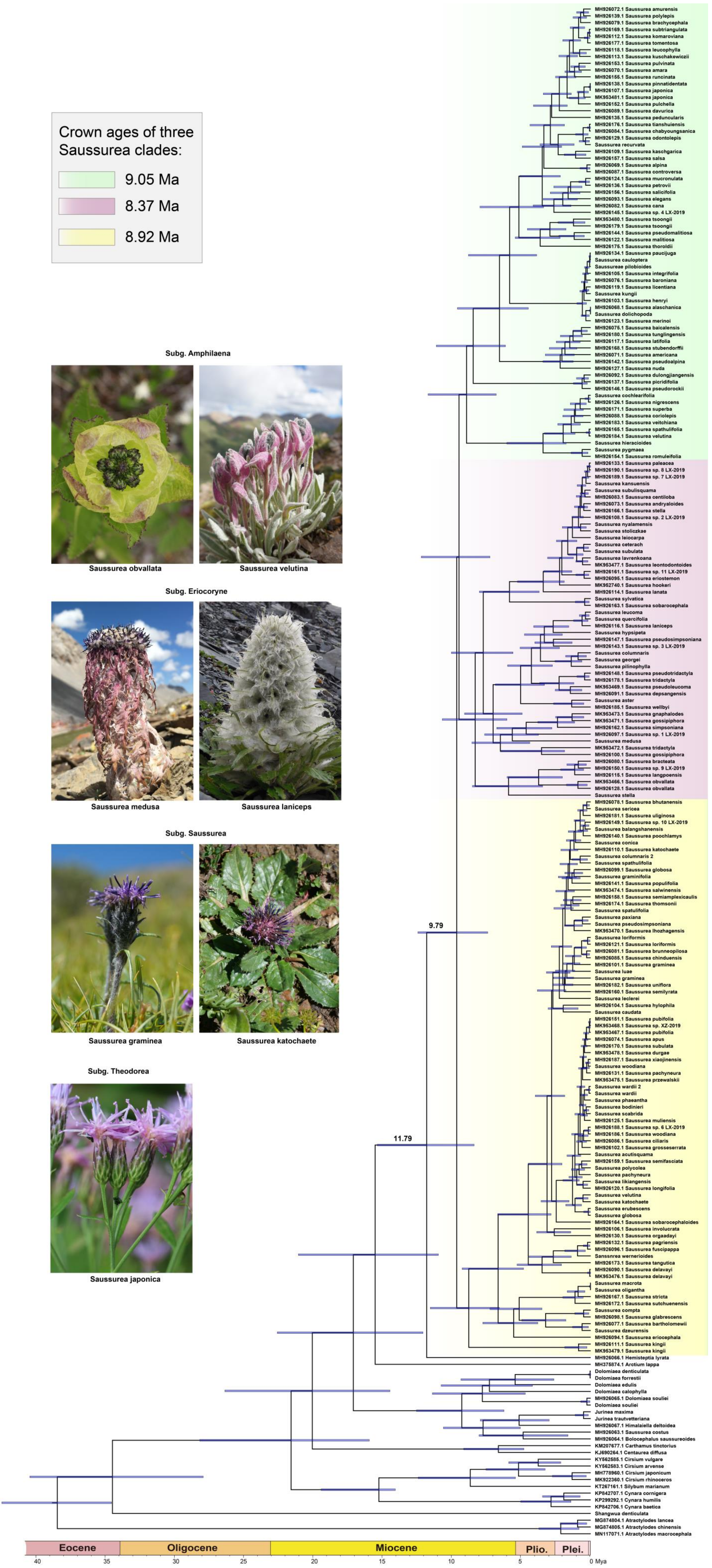

Crown ages of three Saussurea clades:
9.05 Ma
8.37 Ma
8.92 Ma
Subg. Amphilaena
Saussurea obvallata
Saussurea velutina
Subg. Eriocoryne
Saussurea medusa
Saussurea laniceps
Subg. Saussurea
Saussurea graminea
Saussurea katochaete
Subg. Theodorea
Saussurea japonica
9.79
11.79
Eocene
Oligocene
Miocene
Plio.
Plei.
40
35
30
25
20
15
10
5
0 Mya

**Fig. S1** Dated phylogeny of *Saussurea*. The time of three clades beginning to diversify is provided in the box, and the stem and crown ages of *Saussurea* are labeled on the nodes. Node bars provided are 95% height posterior density (HPD) intervals for mean node ages. Representative taxa from four morphological based subgenera are shown.

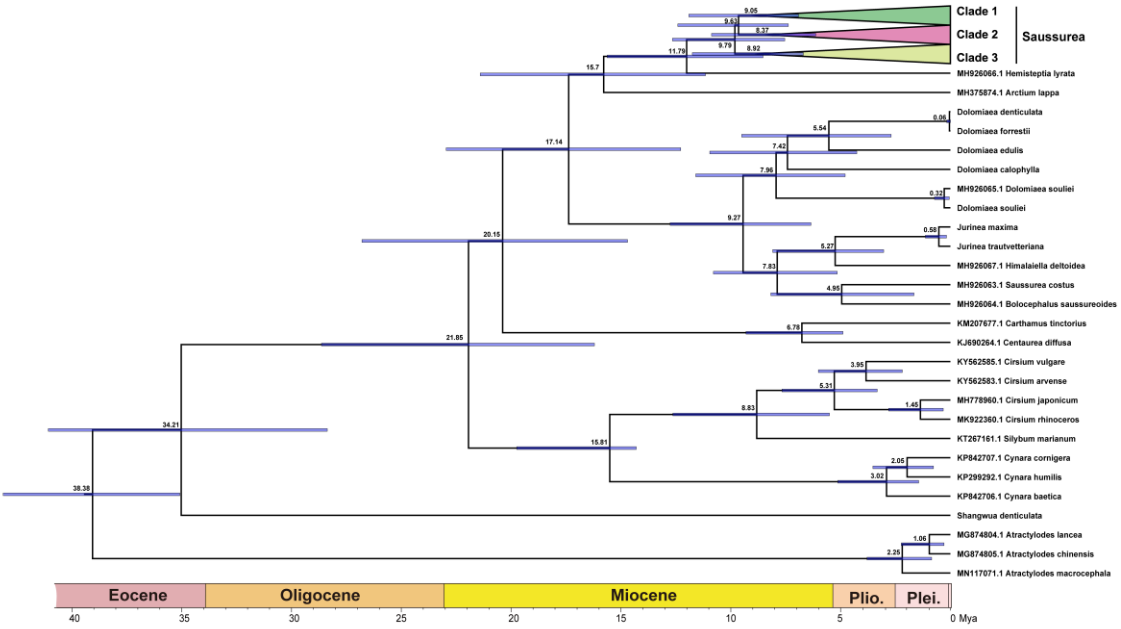

**Fig. S2** Dated phylogeny showing the diversification time of three clades of *Saussurea* is almost simultaneous. Species in the three clades are not shown. Mean node ages are provided and node bars provided are 95% HPD intervals for mean node ages.

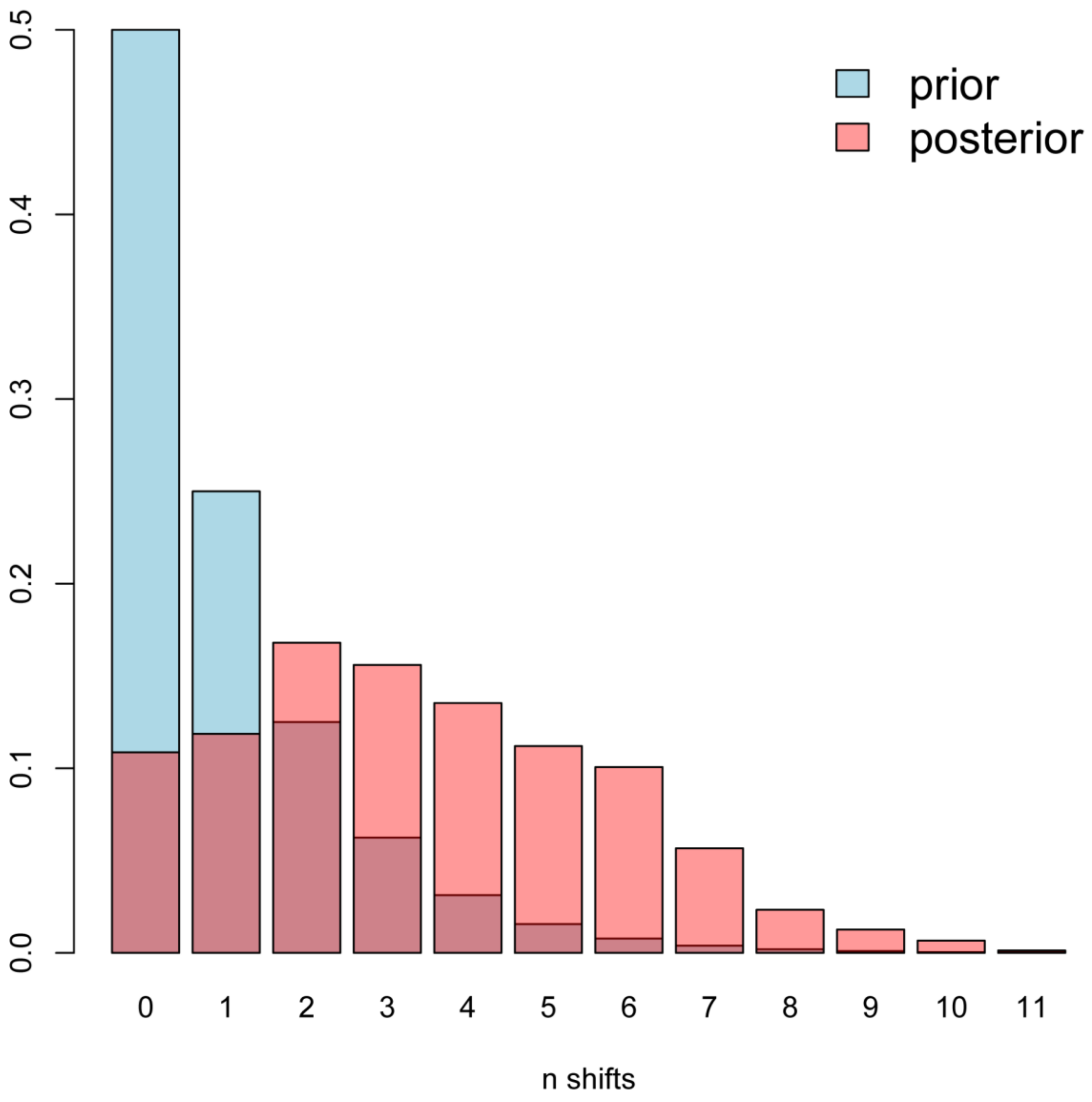

**Fig. S3** Summary of posterior probability of each shift rate in the BAMM analysis.

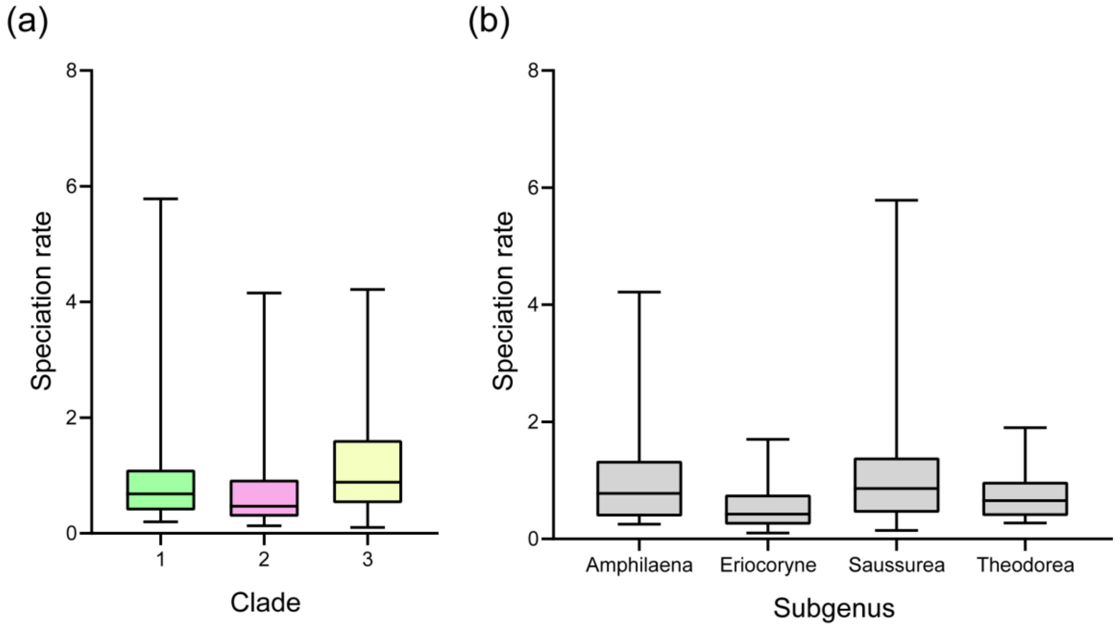

**Fig. S4** Speciation rates estimated by DR statistic (Jetz *et al*. 2012). Tip rates of (a) three clades and (b) four morphology-based subgenera of *Saussurea*.

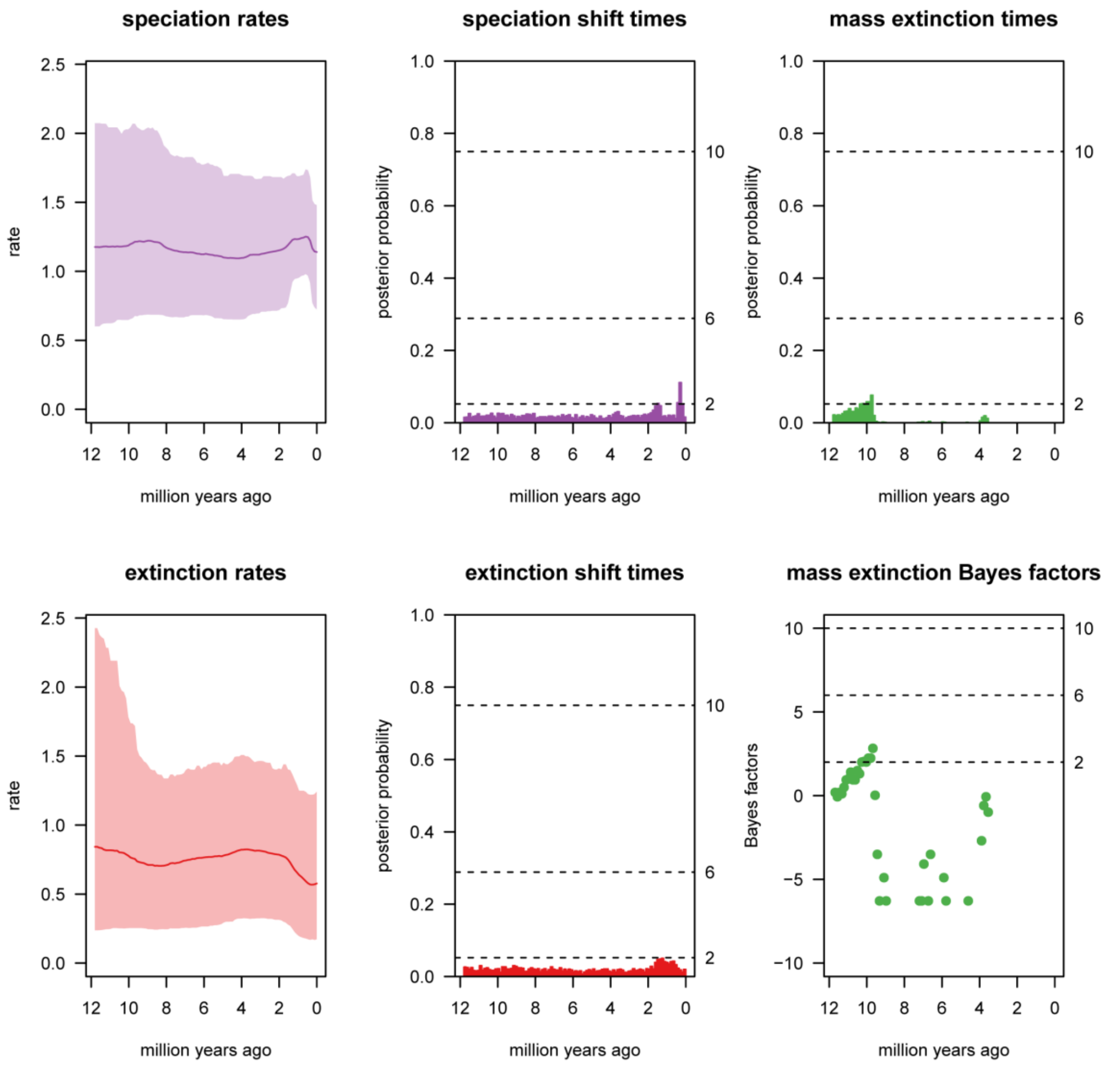

**Fig. S5** Rate-through-time plot from the TESS analysis, showing speciation and exaction shifts have higher posterior probability during the Pleistocene (ca. 2 Mya).

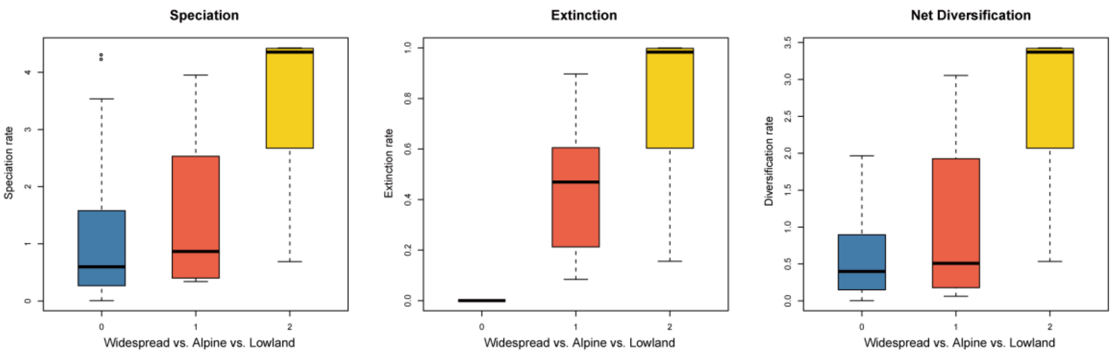

**Fig. S6** Range-dependent diversification of *Saussurea* inferred from GeoHiSSE analysis. Speciation, extinction and net diversification rates of geographical habitats: widespread (0) vs. alpine (1) vs. lowland (2) are provided.

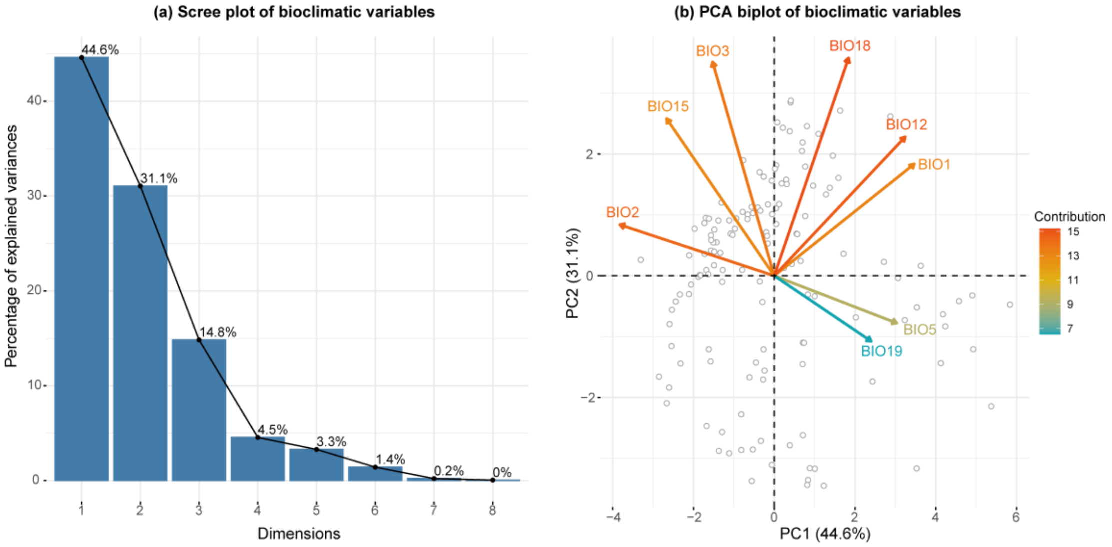

**Fig. S7** PCA results of eight bioclimatic variables representing climate lability of *Saussurea* species. (a) The percentage of explained variance is provided as a scree plot. (b) PCA biplot shows the contribution of eight bioclimatic variables to first two PCs.

**Table S1** Sample information in the present study. -, not applicable.

| Sample names | Voucher specimens | Genebank Accession |
|---|---|---|
| Dolomiaea_calophylla | ZJW5631 | |
| Dolomiaea_denticulata | SunH-07ZX-3416 | |
| Dolomiaea_edulis | ZJW5130 | |
| Dolomiaea_forrestii | FSC-600 | |
| Dolomiaea_souliei | FSC-323 | |
| Jurinea_maxima | Sunhang17405 | |
| Jurinea_trautvetteriana | Sunhang17322 | |
| KJ690264.1_Centaurea_diffusa | - | KJ690264.1 |
| KM207677.1_Carthamus_tinctorius | - | KM207677.1 |
| KP299292.1_Cynara_humilis | - | KP299292.1 |
| KP842706.1_Cynara_baetica | - | KP842706.1 |
| KP842707.1_Cynara_cornigera | - | KP842707.1 |
| KT267161.1_Silybum_marianum | - | KT267161.1 |
| KY562583.1_Cirsium_arvense | - | KY562583.1 |
| KY562585.1_Cirsium_vulgare | - | KY562585.1 |
| MG874804.1_Atractylodes_lancea | - | MG874804.1 |
| MG874805.1_Atractylodes_chinensis | - | MG874805.1 |
| MH375874.1_Arctium_lappa | - | MH375874.1 |
| MH778960.1_Cirsium_japonicum | - | MH778960.1 |
| MH926063.1_Saussurea_costus | - | MH926063.1 |
| MH926064.1_Bolocephalus_saussureoides | - | MH926064.1 |
| MH926065.1_Dolomiaea_souliei | - | MH926065.1 |
| MH926066.1_Hemisteptia_lyrata | - | MH926066.1 |
| MH926067.1_Himalaiella_deltoidea | - | MH926067.1 |
| MH926068.1_Saussurea_alaschanica | - | MH926068.1 |
| MH926069.1_Saussurea_alpina | - | MH926069.1 |
| MH926070.1_Saussurea_amara | - | MH926070.1 |
| MH926071.1_Saussurea_americana | - | MH926071.1 |
| MH926072.1_Saussurea_amurensis | - | MH926072.1 |
| MH926073.1_Saussurea_andryaloides | - | MH926073.1 |
| MH926074.1_Saussurea_apus | - | MH926074.1 |
| MH926075.1_Saussurea_baicalensis | - | MH926075.1 |
| MH926076.1_Saussurea_baroniana | - | MH926076.1 |
| MH926077.1_Saussurea_bartholomewii | - | MH926077.1 |
| MH926078.1_Saussurea_bhutanensis | - | MH926078.1 |
| MH926079.1_Saussurea_brachycephala | - | MH926079.1 |
| MH926080.1_Saussurea_bracteata | - | MH926080.1 |
| MH926081.1_Saussurea_brunneopilosa | - | MH926081.1 |
| MH926082.1_Saussurea_cana | - | MH926082.1 |

| | | |
|---|---|---|
| MH926083.1_Saussurea_centiloba | - | MH926083.1 |
| MH926084.1_Saussurea_chabyoungsanica | - | MH926084.1 |
| MH926085.1_Saussurea_chinduensis | - | MH926085.1 |
| MH926086.1_Saussurea_ciliaris | - | MH926086.1 |
| MH926087.1_Saussurea_controversa | - | MH926087.1 |
| MH926088.1_Saussurea_coriolepis | - | MH926088.1 |
| MH926089.1_Saussurea_davurica | - | MH926089.1 |
| MH926090.1_Saussurea_delavayi | - | MH926090.1 |
| MH926091.1_Saussurea_depsangensis | - | MH926091.1 |
| MH926092.1_Saussurea_dulongjiangensis | - | MH926092.1 |
| MH926093.1_Saussurea_elegans | - | MH926093.1 |
| MH926094.1_Saussurea_eriocephala | - | MH926094.1 |
| MH926095.1_Saussurea_eriostemon | - | MH926095.1 |
| MH926096.1_Saussurea_fuscipappa | - | MH926096.1 |
| MH926097.1_Saussurea_sp._1_LX-2019 | - | MH926097.1 |
| MH926098.1_Saussurea_glabrescens | - | MH926098.1 |
| MH926099.1_Saussurea_globosa | - | MH926099.1 |
| MH926100.1_Saussurea_gossipiphora | - | MH926100.1 |
| MH926101.1_Saussurea_graminea | - | MH926101.1 |
| MH926102.1_Saussurea_grosseserrata | - | MH926102.1 |
| MH926103.1_Saussurea_henryi | - | MH926103.1 |
| MH926104.1_Saussurea_hylophila | - | MH926104.1 |
| MH926105.1_Saussurea_integrifolia | - | MH926105.1 |
| MH926106.1_Saussurea_involucrata | - | MH926106.1 |
| MH926107.1_Saussurea_japonica | - | MH926107.1 |
| MH926108.1_Saussurea_sp._2_LX-2019 | - | MH926108.1 |
| MH926109.1_Saussurea_kaschgarica | - | MH926109.1 |
| MH926110.1_Saussurea_katochaete | - | MH926110.1 |
| MH926111.1_Saussurea_kingii | - | MH926111.1 |
| MH926112.1_Saussurea_komaroviana | - | MH926112.1 |
| MH926113.1_Saussurea_kuschakewiczii | - | MH926113.1 |
| MH926114.1_Saussurea_lanata | - | MH926114.1 |
| MH926115.1_Saussurea_langpoensis | - | MH926115.1 |
| MH926116.1_Saussurea_laniceps | - | MH926116.1 |
| MH926117.1_Saussurea_latifolia | - | MH926117.1 |
| MH926118.1_Saussurea_leucophylla | - | MH926118.1 |
| MH926119.1_Saussurea_licentiana | - | MH926119.1 |
| MH926120.1_Saussurea_longifolia | - | MH926120.1 |
| MH926121.1_Saussurea_loriformis | - | MH926121.1 |
| MH926122.1_Saussurea_malitiosa | - | MH926122.1 |
| MH926123.1_Saussurea_merinoi | - | MH926123.1 |
| MH926124.1_Saussurea_mucronulata | - | MH926124.1 |
| MH926125.1_Saussurea_muliensis | - | MH926125.1 |

| | | |
|---|---|---|
| MH926126.1_Saussurea_nigrescens | - | MH926126.1 |
| MH926127.1_Saussurea_nuda | - | MH926127.1 |
| MH926128.1_Saussurea_obvallata | - | MH926128.1 |
| MH926129.1_Saussurea_odontolepis | - | MH926129.1 |
| MH926130.1_Saussurea_orgaadayi | - | MH926130.1 |
| MH926131.1_Saussurea_pachyneura | - | MH926131.1 |
| MH926132.1_Saussurea_pagriensis | - | MH926132.1 |
| MH926133.1_Saussurea_paleacea | - | MH926133.1 |
| MH926134.1_Saussurea_paucijuga | - | MH926134.1 |
| MH926135.1_Saussurea_peduncularis | - | MH926135.1 |
| MH926136.1_Saussurea_petrovii | - | MH926136.1 |
| MH926137.1_Saussurea_picridifolia | - | MH926137.1 |
| MH926138.1_Saussurea_pinnatidentata | - | MH926138.1 |
| MH926139.1_Saussurea_polylepis | - | MH926139.1 |
| MH926140.1_Saussurea_poochlamys | - | MH926140.1 |
| MH926141.1_Saussurea_populifolia | - | MH926141.1 |
| MH926142.1_Saussurea_pseudoalpina | - | MH926142.1 |
| MH926143.1_Saussurea_sp._3_LX-2019 | - | MH926143.1 |
| MH926144.1_Saussurea_pseudomalitiosa | - | MH926144.1 |
| MH926145.1_Saussurea_sp._4_LX-2019 | - | MH926145.1 |
| MH926146.1_Saussurea_pseudorockii | - | MH926146.1 |
| MH926147.1_Saussurea_pseudosimpsoniana | - | MH926147.1 |
| MH926148.1_Saussurea_pseudotridactyla | - | MH926148.1 |
| MH926149.1_Saussurea_sp._10_LX-2019 | - | MH926149.1 |
| MH926150.1_Saussurea_sp._9_LX-2019 | - | MH926150.1 |
| MH926151.1_Saussurea_pubifolia | - | MH926151.1 |
| MH926152.1_Saussurea_pulchella | - | MH926152.1 |
| MH926153.1_Saussurea_pulvinata | - | MH926153.1 |
| MH926154.1_Saussurea_romuleifolia | - | MH926154.1 |
| MH926155.1_Saussurea_runcinata | - | MH926155.1 |
| MH926156.1_Saussurea_salicifolia | - | MH926156.1 |
| MH926157.1_Saussurea_salsa | - | MH926157.1 |
| MH926158.1_Saussurea_semiamplexicaulis | - | MH926158.1 |
| MH926159.1_Saussurea_semifasciata | - | MH926159.1 |
| MH926160.1_Saussurea_semilyrata | - | MH926160.1 |
| MH926161.1_Saussurea_sp._11_LX-2019 | - | MH926161.1 |
| MH926162.1_Saussurea_simpsoniana | - | MH926162.1 |
| MH926163.1_Saussurea_sobarocephala | - | MH926163.1 |
| MH926164.1_Saussurea_sobarocephaloides | - | MH926164.1 |
| MH926165.1_Saussurea_spathulifolia | - | MH926165.1 |
| MH926166.1_Saussurea_stella | - | MH926166.1 |
| MH926167.1_Saussurea_stricta | - | MH926167.1 |
| MH926168.1_Saussurea_stubendorffii | - | MH926168.1 |

| | | |
|---|---|---|
| MH926169.1_Saussurea_subtriangulata | - | MH926169.1 |
| MH926170.1_Saussurea_subulata | - | MH926170.1 |
| MH926171.1_Saussurea_superba | - | MH926171.1 |
| MH926172.1_Saussurea_sutchuenensis | - | MH926172.1 |
| MH926173.1_Saussurea_tangutica | - | MH926173.1 |
| MH926174.1_Saussurea_thomsonii | - | MH926174.1 |
| MH926175.1_Saussurea_thoroldii | - | MH926175.1 |
| MH926176.1_Saussurea_tianshuiensis | - | MH926176.1 |
| MH926177.1_Saussurea_tomentosa | - | MH926177.1 |
| MH926178.1_Saussurea_tridactyla | - | MH926178.1 |
| MH926179.1_Saussurea_tsoongii | - | MH926179.1 |
| MH926180.1_Saussurea_tunglingensis | - | MH926180.1 |
| MH926181.1_Saussurea_uliginosa | - | MH926181.1 |
| MH926182.1_Saussurea_uniflora | - | MH926182.1 |
| MH926183.1_Saussurea_veitchiana | - | MH926183.1 |
| MH926184.1_Saussurea_velutina | - | MH926184.1 |
| MH926185.1_Saussurea_wellbyi | - | MH926185.1 |
| MH926186.1_Saussurea_woodiana | - | MH926186.1 |
| MH926187.1_Saussurea_xiaojinensis | - | MH926187.1 |
| MH926188.1_Saussurea_sp._6_LX-2019 | - | MH926188.1 |
| MH926189.1_Saussurea_sp._7_LX-2019 | - | MH926189.1 |
| MH926190.1_Saussurea_sp._8_LX-2019 | - | MH926190.1 |
| MK922360.1_Cirsium_rhinoceros | - | MK922360.1 |
| MK952740.1_Saussurea_hookeri | - | MK952740.1 |
| MK953466.1_Saussurea_obvallata | - | MK953466.1 |
| MK953467.1_Saussurea_pubifolia | - | MK953467.1 |
| MK953468.1_Saussurea_sp._XZ-2019 | - | MK953468.1 |
| MK953469.1_Saussurea_pseudoleucoma | - | MK953469.1 |
| MK953470.1_Saussurea_lhozhagensis | - | MK953470.1 |
| MK953471.1_Saussurea_gossipiphora | - | MK953471.1 |
| MK953472.1_Saussurea_tridactyla | - | MK953472.1 |
| MK953473.1_Saussurea_gnaphalodes | - | MK953473.1 |
| MK953474.1_Saussurea_salwinensis | - | MK953474.1 |
| MK953475.1_Saussurea_przewalskii | - | MK953475.1 |
| MK953476.1_Saussurea_delavayi | - | MK953476.1 |
| MK953477.1_Saussurea_leontodontoides | - | MK953477.1 |
| MK953478.1_Saussurea_durgae | - | MK953478.1 |
| MK953479.1_Saussurea_kingii | - | MK953479.1 |
| MK953480.1_Saussurea_tsoongii | - | MK953480.1 |
| MK953481.1_Saussurea_japonica | - | MK953481.1 |
| MN117071.1_Atractylodes_macrocephala | - | MN117071.1 |
| Saussurea_acutisquama | KUN40464 | |
| Saussurea_aster | FSC-81 | |

| | |
|---|---|
| Saussurea_balangshanensis | FSC-176 |
| Saussurea_bodinieri | ZhangDC-07ZX-1812 |
| Saussurea_caudata | KUN38405 |
| Saussurea_cauloptera | YLS2019727 |
| Saussurea_ceterach | ZJW5762 |
| Saussurea_cochlearifolia | SunH-07ZX-1581 |
| Saussurea_columnaris | NK11345 |
| Saussurea_columnaris_2 | XDASDOUT-61 |
| Saussurea_compta | KUN39638 |
| Saussurea_conica | ZhangDC-07ZX-1820 |
| Saussurea_dolichopoda | FSC-238 |
| Saussurea_dzeurensis | KUN34764 |
| Saussurea_erubescens | FSC-425 |
| Saussurea_georgei | SunH-07ZX-3483 |
| Saussurea_globosa | FSC-486 |
| Saussurea_graminea | FSC-51 |
| Saussurea_graminifolia | XU Bo-1005 |
| Saussurea_hieracioides | FSC-719 |
| Saussurea_hypsipeta | FSC-117 |
| Saussurea_kansuensis | KUN36676 |
| Saussurea_katochaete | FSC-612 |
| Saussurea_kungii | YLS2019826 |
| Saussurea_lavrenkoana | KUN36161 |
| Saussurea_leclerei | ZJW5885 |
| Saussurea_leiocarpa | SunH-07ZX-2750 |
| Saussurea_leucoma | FSC-621 |
| Saussurea_likiangensis | SunH-07ZX-2065 |
| Saussurea_loriformis | FSC-434 |
| Saussurea_luae | SunH-07ZX-3437 |
| Saussurea_macrota | YLS2019820 |
| Saussurea_medusa | FSC-030 |
| Saussurea_nyalamensis | ZJW5183 |
| Saussurea_oligantha | YLS2019728 |
| Saussurea_pachyneura | FSC-567 |
| Saussurea_paxiana | Deng5472 |
| Saussurea_phaeantha | FSC-141 |
| Saussurea_pilinophylla | Deng5477 |
| Saussurea_pilobioides | KUN40171 |
| Saussurea_polycolea | FSC-287 |
| Saussurea_pseudosimpsoniana | ZJW6371 |
| Saussurea_pygmaea | SunH-07ZX-1733 |
| Saussurea_quercifolia | FSC-626 |
| Saussurea_recurvata | YLS2019729 |

| | |
|---|---|
| Saussurea_scabrida | SunH-07ZX-3569 |
| Saussurea_sericea | SunH-07ZX-0933 |
| Saussurea_spathulifolia | FSC-616 |
| Saussurea_spatulifolia | KUN33427 |
| Saussurea_stella | FSC-50 |
| Saussurea_stoliczkae | SunH-07ZX-3592 |
| Saussurea_subulata | FSC-418 |
| Saussurea_subulisquama | KUN40458 |
| Saussurea_sylvatica | FSC-281 |
| Saussurea_velutina | FSC-142 |
| Saussurea_wardii | FSC-720 |
| Saussurea_wardii_2 | FSC-721 |
| Saussurea_wernerioides | FSC-630 |
| Saussurea_woodiana | FSC-237 |
| Shangwua_denticulata | ZJW5877 |

**Table S2** Traits coding regime in the present study. Nine characters including four binary morphological traits, four multistate morphological traits and geographical habitats were selected and coded based on descriptions in eFloras (http://www.efloras.org/), herbarium specimens and taxonomic literature, or were manual checked directly using online herbarium specimens from the Chinese Virtual Herbarium (http://www.cvh.ac.cn/), JSTOR (https://plants.jstor.org/), and field collections. -, not available.

| **Species** | **Stemmed (1) vs Stemless (0)** | **Haired (1) vs glabrous (0) stem** | **Leaf entire (0) vs pintely divided (1) vs both (2)** | **Leaves adaxially glabrous (0) vs sparsely haired (1) vs densely haired (2)** | **Capitulum solitary (0) vs numerous (1)** | **Phyllaries rows, <5 (0), 5 (1), 6 (2), >6 (3)** | **Phyllaries glabrous (0) vs sparsely haired (1) vs densely haired (2) vs appendage (3)** | **With (1) vs without (0) bracts** | **Geographical habitats: alpine (1) vs. lowland (2) vs. widespread (0)** | **Subgenus** |
|---|---|---|---|---|---|---|---|---|---|---|
| *Saussurea alaschanica* | 1 | 1 | 0 | 0 | 0 | 1 | 1 | 0 | 2 | S |
| *Saussurea alpina* | 1 | 0 | 0 | 0 | 1 | 0 | 0 | 0 | 0 | S |
| *Saussurea amara* | 1 | 0 | 0 | 1 | 1 | 0 | 3 | 0 | 0 | T |
| *Saussurea americana* | 1 | 0 | 0 | 0 | 1 | 0 | 2 | 0 | 2 | - |
| *Saussurea amurensis* | 1 | 0 | 0 | 0 | 1 | 1 | 2 | 0 | 2 | S |
| *Saussurea andryaloides* | 0 | 1 | 1 | 2 | 0 | 2 | 2 | 0 | 1 | S |
| *Saussurea apus* | 0 | 0 | 1 | 0 | 0 | 0 | 0 | 0 | 1 | S |
| *Saussurea baicalensis* | 1 | 1 | 0 | 2 | 1 | 0 | 1 | 1 | 2 | S |
| *Saussurea baroniana* | 1 | 1 | 0 | 1 | 1 | 3 | 2 | 0 | 2 | S |
| *Saussurea bartholomewii* | 1 | 0 | 1 | 1 | 1 | 2 | 2 | 0 | 2 | S |
| *Saussurea bhutanensis* | 0 | 0 | 0 | 0 | 0 | 2 | 2 | 0 | 1 | S |
| *Saussurea brachycephala* | 1 | 1 | 0 | 1 | 1 | 2 | 1 | 0 | 1 | S |

| | | | | | | | | | | |
|---|---|---|---|---|---|---|---|---|---|---|
| *Saussurea bracteata* | 1 | 0 | 0 | 2 | 0 | 0 | 2 | 1 | 1 | A |
| *Saussurea brunneopilosa* | 1 | 1 | 0 | 0 | 0 | 0 | 2 | 0 | 2 | S |
| *Saussurea cana* | 1 | 1 | 1 | 0 | 1 | 1 | 1 | 0 | 2 | S |
| *Saussurea centiloba* | 1 | 1 | 1 | 1 | 0 | 2 | 2 | 0 | 0 | S |
| *Saussurea chabyoungsanica* | 1 | - | 0 | 2 | 1 | 3 | - | 0 | 2 | - |
| *Saussurea chinduensis* | 1 | 1 | 0 | 0 | 1 | 1 | 2 | 0 | 1 | S |
| *Saussurea ciliaris* | 0 | 0 | 0 | 0 | 0 | 0 | 0 | 0 | 1 | S |
| *Saussurea controversa* | 1 | 0 | 0 | 1 | 1 | 1 | 0 | 0 | 2 | S |
| *Saussurea coriolepis* | 1 | 0 | 2 | 1 | 1 | 1 | 3 | 0 | 1 | S |
| *Saussurea davurica* | 1 | 0 | 2 | 0 | 1 | 3 | 0 | 0 | 0 | S |
| *Saussurea depsangensis* | 0 | 0 | 0 | 1 | 1 | 0 | 2 | 0 | 1 | E |
| *Saussurea dulongjiangensis* | 1 | 1 | 0 | 0 | 1 | 1 | 1 | 0 | 1 | S |
| *Saussurea elegans* | 1 | 1 | 2 | 1 | 1 | 1 | 1 | 0 | 2 | S |
| *Saussurea eriocephala* | 1 | 0 | 2 | 0 | 1 | 1 | 2 | 0 | 2 | S |
| *Saussurea eriostemon* | 1 | 0 | 1 | 0 | 0 | 2 | 0 | 0 | 1 | S |
| *Saussurea fuscipappa* | 1 | 1 | 0 | 1 | 1 | 1 | 1 | 0 | 1 | S |
| *Saussurea glabrescens* | 1 | 1 | 0 | 0 | 1 | 2 | 2 | 0 | 1 | S |
| *Saussurea globosa* | 1 | 1 | 0 | 1 | 1 | 0 | 2 | 1 | 0 | A |
| *Saussurea gossipiphora* | 1 | 1 | 0 | 2 | 1 | 0 | 2 | 0 | 1 | E |
| *Saussurea grosseserrata* | 1 | 1 | 2 | 1 | 0 | 2 | 1 | 0 | 0 | S |
| *Saussurea henryi* | 1 | 0 | 1 | 0 | 0 | 1 | 0 | 0 | 2 | S |
| *Saussurea hylophila* | 1 | 0 | 0 | 1 | 0 | 1 | 0 | 0 | 2 | S |
| *Saussurea integrifolia* | 1 | 0 | 0 | 1 | 1 | 2 | 0 | 0 | 2 | S |
| *Saussurea involucrata* | 1 | 0 | 0 | 0 | 1 | 0 | 1 | 1 | 0 | A |

| | | | | | | | | | |
|---|---|---|---|---|---|---|---|---|---|
| *Saussurea japonica* | 1 | 0 | 2 | 0 | 1 | 2 | 3 | 0 | 0 | T |
| *Saussurea kaschgarica* | 1 | 1 | 1 | 1 | 1 | 1 | 0 | 0 | 2 | S |
| *Saussurea katochaete* | 0 | 0 | 0 | 0 | 0 | 0 | 0 | 0 | 0 | S |
| *Saussurea komaroviana* | 1 | - | 0 | - | 0 | - | - | 0 | 2 | - |
| *Saussurea kuschakewiczii* | 0 | 0 | 0 | 1 | 1 | | 2 | 0 | 0 | S |
| *Saussurea lanata* | 1 | 1 | 2 | 2 | 0 | 0 | 2 | 0 | 1 | S |
| *Saussurea langpoensis* | 1 | 1 | 2 | 0 | 0 | 1 | 2 | 0 | 1 | S |
| *Saussurea laniceps* | 1 | 1 | 0 | 2 | 3 | 0 | 2 | 1 | 1 | E |
| *Saussurea latifolia* | 1 | 0 | 0 | 1 | 1 | 1 | 1 | 0 | 2 | S |
| *Saussurea leucophylla* | 1 | 1 | 0 | 2 | 0 | 0 | 2 | 0 | 0 | S |
| *Saussurea licentiana* | 1 | 0 | 0 | 1 | 1 | 0 | 0 | 0 | 2 | - |
| *Saussurea longifolia* | 1 | 1 | 0 | 2 | 0 | 0 | 2 | 1 | 1 | A |
| *Saussurea malitiosa* | 1 | 1 | 1 | 0 | 1 | 3 | 2 | 0 | 1 | S |
| *Saussurea merinoi* | 1 | 0 | 0 | 0 | 1 | 1 | - | 0 | 2 | S |
| *Saussurea mucronulata* | 1 | 1 | 0 | 2 | 1 | 1 | 0 | 0 | 2 | S |
| *Saussurea muliensis* | 1 | 0 | 0 | 2 | 0 | 2 | 2 | 1 | 1 | A |
| *Saussurea nigrescens* | 1 | 0 | 0 | 0 | 1 | 1 | - | 1 | 2 | A |
| *Saussurea nuda* | 1 | 0 | 0 | - | 1 | 0 | 1 | 0 | 2 | - |
| *Saussurea obvallata* | 1 | 0 | 0 | 1 | 1 | 0 | 1 | 1 | 1 | A |
| *Saussurea odontolepis* | 1 | 0 | 1 | 1 | 1 | 1 | 2 | 0 | 2 | - |
| *Saussurea orgaadayi* | 1 | 0 | 0 | 1 | 1 | 1 | 2 | 1 | 2 | A |
| *Saussurea pachyneura* | 1 | 0 | 1 | 1 | 0 | 2 | 1 | 0 | 1 | S |
| *Saussurea pagriensis* | 0 | 0 | 0 | 1 | 0 | 1 | - | 0 | 2 | S |
| *Saussurea paleacea* | 1 | 1 | 0 | 2 | 0 | 1 | 2 | 0 | 1 | S |

| | | | | | | | | | | |
|---|---|---|---|---|---|---|---|---|---|---|
| *Saussurea paucijuga* | 1 | 1 | 2 | 0 | 0 | 0 | 0 | 0 | 2 | S |
| *Saussurea peduncularis* | 1 | 0 | 1 | 0 | 1 | 1 | 1 | 0 | 2 | S |
| *Saussurea petrovii* | 1 | 1 | 0 | 0 | 0 | 1 | 1 | 0 | 2 | S |
| *Saussurea picridifolia* | 1 | 0 | 1 | 0 | 0 | - | - | 0 | 1 | - |
| *Saussurea pinnatidentata* | 1 | 0 | 2 | 0 | 1 | 1 | 3 | 0 | 2 | T |
| *Saussurea polylepis* | 1 | - | 0 | 0 | 1 | - | - | 0 | 2 | - |
| *Saussurea poochlamys* | 0 | 0 | 0 | 0 | 0 | 0 | 0 | 0 | 0 | S |
| *Saussurea populifolia* | 1 | 0 | 0 | 2 | 0 | 2 | 1 | 0 | 0 | S |
| *Saussurea pseudoalpina* | 1 | 0 | 0 | 0 | 1 | 0 | 1 | 0 | 2 | S |
| *Saussurea pseudomalitiosa* | 1 | 1 | 1 | 1 | 1 | 1 | 3 | 0 | 0 | T |
| *Saussurea pseudorockii* | 0 | 0 | 0 | 0 | 0 | 0 | - | 0 | 1 | S |
| *Saussurea pseudosimpsoniana* | 1 | 1 | 2 | 2 | 1 | 0 | 2 | 0 | 1 | E |
| *Saussurea pseudotridactyla* | 1 | 1 | 0 | 1 | 1 | 0 | 2 | 0 | 1 | E |
| *Saussurea pulchella* | 1 | 0 | 2 | 1 | 1 | 3 | 3 | 0 | 2 | T |
| *Saussurea pulvinata* | 1 | 1 | 0 | 0 | 1 | 0 | 1 | 0 | 1 | S |
| *Saussurea romuleifolia* | 1 | 1 | 0 | 0 | 0 | 1 | - | 0 | 0 | S |
| *Saussurea runcinata* | 1 | 0 | 2 | 0 | 1 | 2 | 3 | 0 | 2 | T |
| *Saussurea salicifolia* | 1 | 1 | 0 | 0 | 1 | 1 | 1 | 0 | 0 | S |
| *Saussurea salsa* | 1 | 1 | 2 | 0 | 1 | 3 | 1 | 0 | 2 | S |
| *Saussurea semiamplexicaulis* | 1 | 1 | 0 | 0 | 1 | 2 | 1 | 0 | 2 | S |
| *Saussurea semifasciata* | 1 | 0 | 0 | 0 | 0 | 0 | 2 | 0 | 1 | S |
| *Saussurea semilyrata* | 1 | 1 | 1 | 1 | 0 | 2 | 1 | 0 | 0 | S |
| *Saussurea simpsoniana* | 1 | 0 | 0 | 2 | 1 | - | - | 0 | 1 | A |
| *Saussurea sobarocephala* | 1 | 1 | 0 | 0 | 1 | 0 | 1 | 0 | 0 | S |

| | | | | | | | | | | |
|---|---|---|---|---|---|---|---|---|---|---|
| *Saussurea sobarocephaloides* | 1 | 1 | 0 | 1 | 1 | 2 | 2 | 0 | 2 | S |
| *Saussurea spathulifolia* | 0 | 0 | 0 | 1 | 0 | 0 | 1 | 0 | 0 | S |
| *Saussurea stella* | 0 | 0 | 0 | 0 | 1 | 1 | 0 | 0 | 0 | E |
| *Saussurea stricta* | 1 | 0 | 0 | 0 | 1 | 1 | 0 | 0 | 2 | S |
| *Saussurea stubendorffii* | 1 | 1 | 1 | 0 | 1 | 1 | 2 | 0 | 2 | - |
| *Saussurea subtriangulata* | 1 | 0 | 0 | 0 | 1 | 3 | 3 | 0 | 2 | S |
| *Saussurea subulata* | 0 | 0 | 0 | 0 | 1 | 0 | 0 | 0 | 1 | S |
| *Saussurea superba* | 1 | 1 | 0 | 0 | 0 | 1 | 0 | 0 | 0 | S |
| *Saussurea sutchuenensis* | 1 | 1 | 0 | 0 | 1 | 3 | 0 | 0 | 1 | S |
| *Saussurea tangutica* | 1 | 1 | 0 | 2 | 0 | 0 | 2 | 1 | 1 | A |
| *Saussurea thomsonii* | 0 | 0 | 0 | 0 | 1 | 0 | 0 | 1 | 1 | E |
| *Saussurea thoroldii* | 0 | 0 | 1 | 0 | 1 | 0 | 0 | 0 | 1 | E |
| *Saussurea tianshuiensis* | 1 | 0 | 1 | 0 | 0 | 2 | 0 | 0 | 2 | S |
| *Saussurea tomentosa* | 1 | 1 | 0 | 0 | 0 | 1 | 2 | 1 | 2 | S |
| *Saussurea tridactyla* | 1 | 1 | 0 | 2 | 1 | 0 | 2 | 0 | 1 | E |
| *Saussurea tsoongii* | 1 | 0 | 2 | 1 | 1 | 1 | 3 | 0 | 0 | T |
| *Saussurea tunglingensis* | 1 | 0 | 0 | 0 | 0 | 3 | 2 | 0 | 2 | S |
| *Saussurea uliginosa* | 1 | 0 | 0 | 0 | 1 | 1 | 0 | 1 | 0 | E |
| *Saussurea uniflora* | 1 | 1 | 0 | 0 | 0 | 0 | 2 | 1 | 0 | A |
| *Saussurea veitchiana* | 1 | 1 | 0 | 1 | 1 | 2 | 1 | 1 | 2 | A |
| *Saussurea velutina* | 1 | 1 | 0 | 2 | 0 | 0 | 2 | 1 | 1 | A |
| *Saussurea wellbyi* | 0 | 0 | 0 | 2 | 1 | 1 | 2 | 0 | 1 | E |
| *Saussurea woodiana* | 1 | 0 | 0 | 1 | 0 | 2 | 1 | 0 | 0 | S |
| *Saussurea xiaojinensis* | 1 | 1 | 0 | 0 | 1 | 2 | 1 | 0 | 2 | S |

| | | | | | | | | | | |
|---|---|---|---|---|---|---|---|---|---|---|
| *Saussurea hookeri* | 1 | 0 | 0 | 2 | 0 | 1 | 2 | 0 | 1 | S |
| *Saussurea obvallata* | 1 | 0 | 0 | 1 | 1 | 0 | 1 | 1 | 0 | A |
| *Saussurea pubifolia* | 1 | 1 | 0 | 0 | 0 | - | - | 1 | 1 | A |
| *Saussurea pseudoleucoma* | 1 | 1 | 1 | 2 | 1 | 0 | 1 | 0 | 1 | E |
| *Saussurea lhozhagensis* | 0 | 1 | 0 | 0 | 1 | 0 | 1 | 0 | 1 | E |
| *Saussurea gossipiphora* | 1 | 1 | 0 | 2 | 1 | 0 | 2 | 0 | 1 | E |
| *Saussurea tridactyla* | 1 | 1 | 0 | 2 | 1 | 0 | 2 | 0 | 1 | E |
| *Saussurea gnaphalodes* | 1 | 1 | 0 | 2 | 1 | 0 | 2 | 0 | 1 | E |
| *Saussurea salwinensis* | 1 | 0 | 1 | 0 | 1 | 0 | 1 | 0 | 1 | S |
| *Saussurea przewalskii* | 1 | 1 | 1 | 0 | 1 | 1 | 1 | 0 | 1 | S |
| *Saussurea delavayi* | 1 | 1 | 0 | 0 | 1 | 1 | 1 | 0 | 2 | E |
| *Saussurea leontodontoides* | 1 | 1 | 1 | 1 | 0 | 1 | 0 | 0 | 1 | S |
| *Saussurea durgae* | 0 | 1 | 0 | 2 | 0 | 1 | 1 | 0 | 1 | S |
| *Saussurea kingii* | 0 | 1 | 1 | 1 | 1 | 0 | 1 | 0 | 0 | E |
| *Saussurea tsoongii* | 1 | 0 | 2 | 1 | 1 | 1 | 3 | 0 | 0 | T |
| *Saussurea japonica* | 1 | 0 | 2 | 0 | 1 | 2 | 3 | 0 | 0 | T |
| *Saussurea acutisquama* | 1 | 1 | 0 | 1 | 1 | 1 | 0 | 1 | 1 | A |
| *Saussurea aster* | 0 | 0 | 0 | 1 | 1 | 0 | 2 | 0 | 1 | E |
| *Saussurea balangshanensis* | 1 | 1 | 1 | - | 1 | 3 | - | 1 | 1 | A |
| *Saussurea bodinieri* | 1 | 1 | 1 | 0 | 0 | 2 | 1 | 1 | 0 | S |
| *Saussurea caudata* | 1 | 0 | 0 | 0 | 0 | 1 | 0 | 0 | 0 | S |
| *Saussurea cauloptera* | 1 | 0 | 0 | 0 | 1 | 1 | 1 | 0 | 2 | S |
| *Saussurea ceterach* | 0 | 0 | 1 | 1 | 0 | 3 | 1 | 0 | 1 | S |
| *Saussurea cochlearifolia* | 1 | 0 | 0 | 1 | 0 | 0 | 1 | 0 | 1 | S |

| | | | | | | | | | | |
|---|---|---|---|---|---|---|---|---|---|---|
| *Saussurea columnaris* | 0 | 0 | 0 | 2 | 0 | 1 | 2 | 0 | 1 | S |
| *Saussurea compta* | 1 | 1 | 1 | 1 | 0 | 2 | 2 | 0 | 2 | S |
| *Saussurea conica* | 1 | 1 | 0 | 1 | 1 | - | 2 | 1 | 1 | A |
| *Saussurea dolichopoda* | 1 | 0 | 0 | 0 | 1 | 1 | 0 | 0 | 2 | S |
| *Saussurea dzeurensis* | 1 | 1 | 2 | 1 | 1 | 2 | 1 | 0 | 1 | S |
| *Saussurea erubescens* | 1 | 1 | 0 | 2 | 1 | 2 | 2 | 1 | 0 | A |
| *Saussurea georgei* | 0 | 0 | 2 | 2 | 1 | 0 | 0 | 0 | 1 | E |
| *Saussurea globosa* | 1 | 1 | 0 | 1 | 1 | 0 | 2 | 1 | 0 | A |
| *Saussurea graminea* | 1 | 1 | 0 | 1 | 0 | 1 | 2 | 0 | 1 | S |
| *Saussurea graminifolia* | 1 | 1 | 0 | 0 | 0 | 1 | 2 | 0 | 1 | S |
| *Saussurea hieracioides* | 1 | 1 | 0 | 1 | 0 | 1 | 2 | 0 | 1 | S |
| *Saussurea hypsipeta* | 1 | 1 | 2 | 2 | 1 | 3 | 2 | 0 | 1 | E |
| *Saussurea kansuensis* | 0 | 0 | 1 | 1 | 0 | 0 | 0 | 0 | 2 | S |
| *Saussurea katochaete* | 0 | 0 | 0 | 0 | 0 | 0 | 0 | 0 | 0 | S |
| *Saussurea kungii* | 1 | 0 | 1 | 0 | 0 | 1 | 1 | 1 | 2 | S |
| *Saussurea lavrenkoana* | 0 | 0 | 1 | 0 | 0 | 1 | 0 | 0 | 1 | S |
| *Saussurea leclerei* | 1 | 0 | 0 | 0 | 1 | 1 | 0 | 0 | 2 | S |
| *Saussurea leiocarpa* | 0 | 0 | 1 | 1 | 0 | 1 | 1 | 0 | 1 | S |
| *Saussurea leucoma* | 1 | 1 | 1 | 2 | 1 | 0 | 2 | 0 | 1 | E |
| *Saussurea likiangensis* | 1 | - | 1 | 1 | 1 | - | - | 0 | 0 | - |
| *Saussurea loriformis* | 1 | 1 | 0 | 2 | 0 | 1 | 2 | 0 | 1 | S |
| *Saussurea luae* | 1 | 0 | 0 | 0 | 1 | 2 | 2 | 1 | 1 | A |
| *Saussurea macrota* | 1 | 0 | 0 | 1 | 1 | 2 | 0 | 0 | 2 | S |
| *Saussurea medusa* | 1 | 1 | 0 | 2 | 1 | 3 | 2 | 0 | 1 | E |

| | | | | | | | | | | |
|---|---|---|---|---|---|---|---|---|---|---|
| *Saussurea nyalamensis* | 0 | 0 | 1 | 1 | 0 | 0 | 0 | 0 | 1 | S |
| *Saussurea oligantha* | 1 | 0 | 0 | 1 | 1 | 2 | 3 | 0 | 2 | S |
| *Saussurea pachyneura* | 1 | 0 | 1 | 1 | 0 | 2 | 1 | 0 | 0 | S |
| *Saussurea paxiana* | 1 | 1 | 0 | 0 | 0 | 0 | 2 | 0 | 1 | E |
| *Saussurea phaeantha* | 1 | 1 | 0 | 2 | 1 | 0 | 2 | 1 | 1 | A |
| *Saussurea pilinophylla* | 1 | 0 | 0 | 2 | 0 | 1 | 2 | 1 | 1 | S |
| *Saussurea pilobioides* | 1 | 0 | 0 | 1 | 1 | 1 | 3 | 0 | 0 | S |
| *Saussurea polycolea* | 1 | 1 | 0 | 2 | 0 | 1 | 2 | 1 | 1 | A |
| *Saussurea pseudosimpsoniana* | 1 | 1 | 2 | 2 | 1 | 0 | 2 | 0 | 1 | E |
| *Saussurea pygmaea* | 1 | 1 | 0 | 0 | 0 | - | 2 | 0 | 1 | - |
| *Saussurea quercifolia* | 1 | 1 | 0 | 1 | 1 | 0 | 2 | 0 | 1 | E |
| *Saussurea recurvata* | 1 | 0 | 2 | 1 | 1 | 2 | 0 | 0 | 2 | S |
| *Saussurea scabrida* | 1 | 1 | 2 | 2 | 0 | 1 | 2 | 0 | 1 | S |
| *Saussurea sericea* | 0 | 1 | 0 | 2 | 0 | 0 | 2 | 0 | 1 | S |
| *Saussurea spathulifolia* | 0 | 0 | 0 | 1 | 0 | 0 | 1 | 0 | 1 | S |
| *Saussurea spatulifolia* | 0 | 0 | 0 | 1 | 0 | 0 | 1 | 0 | 1 | S |
| *Saussurea stella* | 0 | 0 | 0 | 0 | 1 | 1 | 0 | 0 | 0 | E |
| *Saussurea stoliczkae* | 0 | 1 | 0 | 1 | 0 | 1 | 1 | 0 | 0 | S |
| *Saussurea subulata* | 0 | 0 | 0 | 0 | 1 | 0 | 0 | 0 | 1 | S |
| *Saussurea subulisquama* | 1 | 1 | 1 | 0 | 0 | 2 | 0 | 0 | 2 | S |
| *Saussurea sylvatica* | 1 | 1 | 0 | 0 | 1 | 1 | 0 | 0 | 0 | S |
| *Saussurea velutina* | 1 | 1 | 0 | 2 | 0 | 0 | 2 | 1 | 1 | A |
| *Saussurea wardii* | 1 | 0 | 1 | 0 | 0 | 2 | 2 | 0 | 1 | S |
| *Saussurea wernerioides* | 0 | 0 | 1 | 0 | 0 | 3 | 0 | 0 | 1 | S |

**Table S3** HiSSE models used in the present study. τ net turnover, ε extinction fraction. Detailed explanation of these models as well as their relationship to one another are described in Beaulieu & O'Meara (2016).

| Model number (#) | Model name |
|---|---|
| 1 | HiSSE full modεl |
| 2 | BiSSE modεl: All fτee |
| 3 | BiSSE modεl: ε0=ε1 |
| 4 | BiSSE modεl: q's equal |
| 5 | BiSSE modεl: q's equal, ε0=ε1 |
| 6 | CID-2: q's equal |
| 7 | CID-2: q's equal and ε's equal |
| 8 | CID-4: q's equal |
| 9 | CID-4: ε's and q's equal |
| 10 | HiSSE: q's equal |
| 11 | HiSSE: q's and ε's equal |
| 12 | HiSSE: τ0A=τ1A=τ0B, ε0A=ε1A=ε1B, q's equal |
| 13 | HiSSE: τ0A=τ1A=τ0B, ε's and q's equal |
| 14 | HiSSE: τ0A=τ0B, ε0A=ε0B |
| 15 | HiSSE: τ0A=τ0B, ε's and q's equal |
| 16 | HiSSE: τ0A=τ1A, ε0A=ε1A, q's equal |
| 17 | HiSSE: τ0A=τ1A, ε's and q's equal |
| 18 | HiSEE: q0B1B=0, q1B0B=0, All other q's equal |
| 19 | HiSSE: ε's equal, q0B1B=0, q1B0B=0, All other q's equal |
| 20 | HiSSE: τ0A=τ1A=τ0B, ε0A=ε1A=ε0B, q0B1B=0, q1B0B=0, All other q's equAl |
| 21 | HiSSE: τ0A=τ1A=τ0B, ε's equal, q0B1B=0, q1B0B=0, All other q's equAl |
| 22 | HiSSE: τ0A=τ0B, ε0A=ε0B, q0B1B=0, q1B0B=0, All other q's equal |
| 23 | HiSSE: τ0A=τ0B, ε's equal, q0B1B=0, q1B0B=0, All other q's equal |
| 24 | HiSSE: τ0A=τ1A, ε0A=ε1A, q0B1B=0, q1B0B=0, All other q's equal |
| 25 | HiSSE: τ0A=τ1A, ε's equal, q0B1B=0, q1B0B=0, All other q's equal |

**Table S4** Speciation rates of *Saussurea* species estimated from BAMM and DR statistic.

| **Species** | **Clade** | **Subgenus** | **BAMM tip rates** | **DR statistic** |
|---|---|---|---|---|
| *Saussurea alaschanica* | 1 | Saussurea | 1.7178 | 0.9785 |
| *Saussurea alpina* | 1 | Saussurea | 0.5162 | 0.4624 |
| *Saussurea amara* | 1 | Theodorea | 0.5529 | 0.8487 |
| *Saussurea americana* | 1 | Saussurea | 0.4748 | 0.3715 |
| *Saussurea amurensis* | 1 | Saussurea | 0.5563 | 1.3477 |
| *Saussurea andryaloides* | 2 | Saussurea | 0.7344 | 1.4670 |
| *Saussurea apus* | 3 | Saussurea | 1.4733 | 2.6409 |
| *Saussurea baicalensis* | 1 | Saussurea | 0.4748 | 0.8506 |
| *Saussurea baroniana* | 1 | Saussurea | 1.7178 | 1.2427 |
| *Saussurea bartholomewii* | 3 | Saussurea | 0.4700 | 0.3813 |
| *Saussurea bhutanensis* | 3 | Saussurea | 0.8338 | 1.8957 |
| *Saussurea brachycephala* | 1 | Saussurea | 0.5563 | 1.0277 |
| *Saussurea bracteata* | 2 | Amphilaena | 0.4699 | 0.4657 |
| *Saussurea brunneopilosa* | 3 | Saussurea | 0.8278 | 1.5289 |
| *Saussurea cana* | 1 | Saussurea | 0.5111 | 0.3509 |
| *Saussurea centiloba* | 2 | Saussurea | 0.7344 | 1.7288 |
| *Saussurea chabyoungsanica* | 1 | Saussurea | 0.5416 | 1.1026 |
| *Saussurea chinduensis* | 3 | Saussurea | 0.8306 | 1.5289 |
| *Saussurea ciliaris* | 3 | Saussurea | 1.4772 | 1.4917 |
| *Saussurea controversa* | 1 | Saussurea | 0.5162 | 0.4624 |
| *Saussurea coriolepis* | 1 | Saussurea | 0.4728 | 0.6460 |
| *Saussurea davurica* | 1 | Saussurea | 0.5383 | 0.3703 |
| *Saussurea depsangensis* | 2 | Eriocoryne | 0.4692 | 0.3796 |
| *Saussurea dulongjiangensis* | 1 | Saussurea | 0.4469 | 0.3789 |
| *Saussurea elegans* | 1 | Saussurea | 0.5160 | 0.4472 |
| *Saussurea eriocephala* | 3 | Saussurea | 0.4301 | 0.1455 |
| *Saussurea eriostemon* | 2 | Saussurea | 0.6439 | 0.5891 |
| *Saussurea fuscipappa* | 3 | Saussurea | 0.4995 | 0.7679 |
| *Saussurea sp. 1 LX-2019* | 2 | Saussurea | 0.4647 | 0.1814 |
| *Saussurea glabrescens* | 3 | Saussurea | 0.4637 | 0.3895 |
| *Saussurea globosa* | 3 | Amphilaena | 0.8284 | 0.7406 |
| *Saussurea gossipiphora* | 2 | Eriocoryne | 0.4532 | 0.1876 |
| *Saussurea sp. 2 XZ* | 3 | Eriocoryne | 0.8274 | 0.7264 |
| *Saussurea grosseserrata* | 3 | Saussurea | 1.4772 | 1.2131 |
| *Saussurea henryi* | 1 | Saussurea | 1.7151 | 0.4777 |
| *Saussurea hylophila* | 3 | Saussurea | 0.8061 | 0.3828 |

| | | | | |
|---|---|---|---|---|
| *Saussurea integrifolia* | 1 | Saussurea | 1.7178 | 1.8977 |
| *Saussurea involucrata* | 3 | Amphilaena | 0.6420 | 0.2716 |
| *Saussurea japonica* | 1 | Theodorea | 0.5630 | 1.9003 |
| *Saussurea sp. 2 LX-2019* | 2 | Theodorea | 0.7325 | 0.9497 |
| *Saussurea kaschgarica* | 1 | Saussurea | 0.5335 | 0.5004 |
| *Saussurea katochaete* | 3 | Saussurea | 0.8225 | 0.7235 |
| *Saussurea komaroviana* | 1 | Saussurea | 0.5610 | 2.4601 |
| *Saussurea kuschakewiczii* | 1 | Saussurea | 0.5555 | 0.8481 |
| *Saussurea lanata* | 2 | Saussurea | 0.4880 | 0.1892 |
| *Saussurea langpoensis* | 2 | Saussurea | 0.4692 | 0.2940 |
| *Saussurea laniceps* | 2 | Eriocoryne | 0.4721 | 0.5096 |
| *Saussurea latifolia* | 1 | Saussurea | 0.4730 | 0.5932 |
| *Saussurea leucophylla* | 1 | Saussurea | 0.5550 | 0.8481 |
| *Saussurea licentiana* | 1 | Saussurea | 1.7179 | 1.2681 |
| *Saussurea longifolia* | 3 | Amphilaena | 1.4355 | 1.0581 |
| *Saussurea malitiosa* | 1 | Saussurea | 0.4783 | 0.3952 |
| *Saussurea merinoi* | 1 | Saussurea | 1.7166 | 0.4937 |
| *Saussurea mucronulata* | 1 | Saussurea | 0.5172 | 0.9072 |
| *Saussurea muliensis* | 3 | Amphilaena | 1.4772 | 1.7354 |
| *Saussurea nigrescens* | 1 | Amphilaena | 0.4728 | 1.5237 |
| *Saussurea nuda* | 1 | Saussurea | 0.4788 | 0.2006 |
| *Saussurea obvallata* | 2 | Amphilaena | 0.4738 | 0.3043 |
| *Saussurea odontolepis* | 1 | Saussurea | 0.5393 | 0.9508 |
| *Saussurea orgaadayi* | 3 | Amphilaena | 0.6538 | 0.2716 |
| *Saussurea pachyneura* | 3 | Saussurea | 1.4772 | 2.1933 |
| *Saussurea pagriensis* | 3 | Saussurea | 0.4995 | 0.7679 |
| *Saussurea paleacea* | 2 | Saussurea | 0.7344 | 4.1539 |
| *Saussurea paucijuga* | 1 | Saussurea | 1.7178 | 5.7834 |
| *Saussurea peduncularis* | 1 | Saussurea | 0.5347 | 0.3182 |
| *Saussurea petrovii* | 1 | Saussurea | 0.5172 | 0.9072 |
| *Saussurea picridifolia* | 1 | Saussurea | 0.4456 | 0.3789 |
| *Saussurea pinnatidentata* | 1 | Theodorea | 0.5630 | 1.9003 |
| *Saussurea polylepis* | 1 | Saussurea | 0.5563 | 1.3477 |
| *Saussurea poochlamys* | 3 | Saussurea | 0.8328 | 1.3039 |
| *Saussurea populifolia* | 3 | Saussurea | 0.8298 | 0.6663 |
| *Saussurea pseudoalpina* | 1 | Saussurea | 0.4723 | 0.2834 |
| *Saussurea sp. 3 LX-2019* | 2 | Saussurea | 0.4710 | 0.3035 |
| *Saussurea pseudomalitiosa* | 1 | Theodorea | 0.4827 | 0.3952 |
| *Saussurea sp. 4 LX-2019* | 1 | Theodorea | 0.5171 | 0.2708 |
| *Saussurea pseudorockii* | 1 | Saussurea | 0.4474 | 0.2083 |
| *Saussurea pseudosimpsoniana* | 2 | Eriocoryne | 0.4663 | 0.3035 |

| | | | | |
|---|---|---|---|---|
| *Saussurea pseudotridactyla* | 2 | Eriocoryne | 0.4703 | 0.4713 |
| *Saussurea sp. 10 LX-2019* | 3 | Eriocoryne | 0.8325 | 1.6996 |
| *Saussurea sp. 9 LX-2019* | 2 | Eriocoryne | 0.4672 | 0.4657 |
| *Saussurea pulchella* | 1 | Theodorea | 0.5571 | 0.6490 |
| *Saussurea pulvinata* | 1 | Saussurea | 0.5506 | 0.8487 |
| *Saussurea romuleifolia* | 1 | Saussurea | 0.4678 | 0.2822 |
| *Saussurea runcinata* | 1 | Theodorea | 0.5574 | 0.6597 |
| *Saussurea salicifolia* | 1 | Saussurea | 0.5172 | 0.5431 |
| *Saussurea salsa* | 1 | Saussurea | 0.5346 | 0.5004 |
| *Saussurea semiamplexicaulis* | 3 | Saussurea | 0.8292 | 0.6747 |
| *Saussurea semifasciata* | 3 | Saussurea | 1.4375 | 1.1502 |
| *Saussurea semilyrata* | 3 | Saussurea | 0.8311 | 0.4669 |
| *Saussurea sp. 11 LX-2019* | 2 | Saussurea | 0.6439 | 0.5891 |
| *Saussurea simpsoniana* | 2 | Amphilaena | 0.4681 | 0.2531 |
| *Saussurea sobarocephala* | 2 | Saussurea | 0.4623 | 0.2636 |
| *Saussurea sobarocephaloides* | 3 | Saussurea | 0.7994 | 0.3196 |
| *Saussurea spathulifolia* | 1 | Saussurea | 0.4726 | 0.6835 |
| *Saussurea stella* | 2 | Eriocoryne | 0.7344 | 1.4670 |
| *Saussurea stricta* | 3 | Saussurea | 0.4713 | 0.4465 |
| *Saussurea stubendorffii* | 1 | Saussurea | 0.4741 | 0.4766 |
| *Saussurea subtriangulata* | 1 | Saussurea | 0.5604 | 2.4601 |
| *Saussurea subulata* | 3 | Saussurea | 1.4772 | 2.2124 |
| *Saussurea superba* | 1 | Saussurea | 0.4728 | 0.8674 |
| *Saussurea sutchuenensis* | 3 | Saussurea | 0.4675 | 0.2802 |
| *Saussurea tangutica* | 3 | Amphilaena | 0.4864 | 0.2627 |
| *Saussurea thomsonii* | 3 | Eriocoryne | 0.8293 | 0.6747 |
| *Saussurea thoroldii* | 1 | Eriocoryne | 0.4726 | 0.2079 |
| *Saussurea tianshuiensis* | 1 | Saussurea | 0.5416 | 1.1026 |
| *Saussurea tomentosa* | 1 | Saussurea | 0.5604 | 1.3937 |
| *Saussurea tridactyla* | 2 | Eriocoryne | 0.4703 | 0.4713 |
| *Saussurea tsoongii* | 1 | Theodorea | 0.4805 | 0.3912 |
| *Saussurea tunglingensis* | 1 | Saussurea | 0.4748 | 0.8506 |
| *Saussurea uliginosa* | 3 | Eriocoryne | 0.8338 | 1.2972 |
| *Saussurea uniflora* | 3 | Amphilaena | 0.8223 | 0.5027 |
| *Saussurea veitchiana* | 1 | Amphilaena | 0.4728 | 0.5017 |
| *Saussurea velutina* | 1 | Amphilaena | 0.4726 | 0.6835 |
| *Saussurea wellbyi* | 2 | Eriocoryne | 0.4660 | 0.2577 |
| *Saussurea woodiana* | 3 | Saussurea | 1.4772 | 2.3801 |
| *Saussurea xiaojinensis* | 3 | Saussurea | 1.4772 | 3.1767 |
| *Saussurea sp. 6 LX-2019* | 3 | Saussurea | 1.4772 | 2.3801 |
| *Saussurea sp. 7 LX-2019* | 2 | Saussurea | 0.7344 | 2.6078 |

| | | | | |
|---|---|---|---|---|
| *Saussurea sp. 8 LX-2019* | 2 | Saussurea | 0.7344 | 4.1539 |
| *Saussurea hookeri* | 2 | Saussurea | 0.6075 | 0.2657 |
| *Saussurea obvallata* | 2 | Amphilaena | 0.4687 | 0.3043 |
| *Saussurea pubifolia* | 3 | Amphilaena | 1.4772 | 4.2175 |
| *Saussurea sp. XZ-2019* | 3 | Amphilaena | 1.4772 | 4.2175 |
| *Saussurea pseudoleucoma* | 2 | Eriocoryne | 0.4705 | 0.3796 |
| *Saussurea lhozhagensis* | 3 | Eriocoryne | 0.8301 | 0.6045 |
| *Saussurea gossipiphora* | 2 | Eriocoryne | 0.4704 | 0.3777 |
| *Saussurea tridactyla* | 2 | Eriocoryne | 0.4572 | 0.1876 |
| *Saussurea gnaphalodes* | 2 | Eriocoryne | 0.4704 | 0.3777 |
| *Saussurea salwinensis* | 3 | Saussurea | 0.8291 | 0.5618 |
| *Saussurea przewalskii* | 3 | Saussurea | 1.4736 | 1.3913 |
| *Saussurea delavayi* | 3 | Eriocoryne | 0.4869 | 0.2031 |
| *Saussurea leontodontoides* | 2 | Saussurea | 0.6998 | 0.8940 |
| *Saussurea durgae* | 3 | Saussurea | 1.4772 | 1.8677 |
| *Saussurea kingii* | 3 | Eriocoryne | 0.4204 | 0.1026 |
| *Saussurea tsoongii* | 1 | Theodorea | 0.4779 | 0.3912 |
| *Saussurea japonica* | 1 | Theodorea | 0.5617 | 0.9874 |
| *Saussurea acutisquama* | 3 | Amphilaena | 1.4675 | 0.7781 |
| *Saussurea aster* | 2 | Eriocoryne | 0.4679 | 0.2577 |
| *Saussurea balangshanensis* | 3 | Amphilaena | 0.8322 | 1.6996 |
| *Saussurea bodinieri* | 3 | Saussurea | 1.4772 | 2.2266 |
| *Saussurea caudata* | 3 | Saussurea | 0.8033 | 0.3828 |
| *Saussurea cauloptera* | 1 | Saussurea | 1.7178 | 5.7834 |
| *Saussurea ceterach* | 2 | Saussurea | 0.7029 | 0.9298 |
| *Saussurea cochlearifolia* | 1 | Saussurea | 0.4728 | 1.5237 |
| *Saussurea columnaris* | 2 | Saussurea | 0.4700 | 0.5201 |
| *Saussurea compta* | 3 | Saussurea | 0.4674 | 0.3895 |
| *Saussurea conica* | 3 | Amphilaena | 0.8320 | 0.8070 |
| *Saussurea dolichopoda* | 1 | Saussurea | 1.7178 | 0.9785 |
| *Saussurea dzeurensis* | 3 | Saussurea | 0.4699 | 0.3813 |
| *Saussurea erubescens* | 3 | Amphilaena | 1.3900 | 1.0832 |
| *Saussurea georgei* | 2 | Eriocoryne | 0.4700 | 0.5201 |
| *Saussurea globosa* | 3 | Amphilaena | 1.3900 | 1.0832 |
| *Saussurea graminea* | 3 | Saussurea | 0.8254 | 0.5400 |
| *Saussurea graminifolia* | 3 | Saussurea | 0.8330 | 0.7406 |
| *Saussurea hieracioides* | 1 | Saussurea | 0.4714 | 0.2366 |
| *Saussurea hypsipeta* | 2 | Eriocoryne | 0.4726 | 0.3028 |
| *Saussurea kansuensis* | 2 | Saussurea | 0.7344 | 1.8547 |
| *Saussurea katochaete* | 3 | Saussurea | 1.3899 | 0.8971 |
| *Saussurea kungii* | 1 | Saussurea | 1.7179 | 1.2681 |

| | | | | |
|---|---|---|---|---|
| *Saussurea lavrenkoana* | 2 | Saussurea | 0.6979 | 0.8940 |
| *Saussurea leclerei* | 3 | Saussurea | 0.7962 | 0.3632 |
| *Saussurea leiocarpa* | 2 | Saussurea | 0.7093 | 0.5551 |
| *Saussurea leucoma* | 2 | Eriocoryne | 0.4721 | 0.8537 |
| *Saussurea likiangensis* | 3 | Saussurea | 1.4355 | 1.0581 |
| *Saussurea loriformis* | 3 | Saussurea | 0.8301 | 0.9841 |
| *Saussurea luae* | 3 | Amphilaena | 0.8201 | 0.6362 |
| *Saussurea macrota* | 3 | Saussurea | 0.4736 | 0.8856 |
| *Saussurea medusa* | 2 | Eriocoryne | 0.4573 | 0.1567 |
| *Saussurea nyalamensis* | 2 | Saussurea | 0.7288 | 0.9654 |
| *Saussurea oligantha* | 3 | Saussurea | 0.4736 | 0.8856 |
| *Saussurea pachyneura* | 3 | Saussurea | 1.4202 | 0.9180 |
| *Saussurea paxiana* | 3 | Eriocoryne | 0.8291 | 0.8889 |
| *Saussurea phaeantha* | 3 | Amphilaena | 1.4767 | 1.7322 |
| *Saussurea pilinophylla* | 2 | Saussurea | 0.4716 | 0.3407 |
| *Saussurea pilobioides* | 1 | Saussurea | 1.7178 | 3.3213 |
| *Saussurea polycolea* | 3 | Amphilaena | 1.4375 | 1.1502 |
| *Saussurea pseudosimpsoniana* | 3 | Eriocoryne | 0.8291 | 0.8889 |
| *Saussurea pygmaea* | 1 | Saussurea | 0.4684 | 0.2822 |
| *Saussurea quercifolia* | 2 | Eriocoryne | 0.4716 | 0.8537 |
| *Saussurea recurvata* | 1 | Saussurea | 0.5393 | 0.9508 |
| *Saussurea scabrida* | 3 | Saussurea | 1.4772 | 2.2266 |
| *Saussurea sericea* | 3 | Saussurea | 0.8338 | 1.8957 |
| *Saussurea sp. 1 XZ* | 3 | Saussurea | 0.8276 | 0.8632 |
| *Saussurea sp. 3 XZ* | 3 | Saussurea | 1.4772 | 3.1767 |
| *Saussurea spathulifolia* | 3 | Saussurea | 0.8295 | 0.8632 |
| *Saussurea spatulifolia* | 3 | Saussurea | 0.8296 | 0.5964 |
| *Saussurea stella* | 2 | Eriocoryne | 0.4520 | 0.1323 |
| *Saussurea stoliczkae* | 2 | Saussurea | 0.7288 | 0.9654 |
| *Saussurea subulata* | 2 | Saussurea | 0.7029 | 0.9298 |
| *Saussurea subulisquama* | 2 | Saussurea | 0.7344 | 1.7288 |
| *Saussurea sylvatica* | 2 | Saussurea | 0.4623 | 0.2636 |
| *Saussurea velutina* | 3 | Amphilaena | 1.3900 | 0.8971 |
| *Saussurea wardii* | 3 | Saussurea | 1.4772 | 1.7322 |
| *Saussurea wernerioides* | 3 | Theodorea | 0.4931 | 0.4203 |

**Table S5** Statistic summary of *Saussurea* speciation rates from BAMM analysis. ANOVA analysis shows the comparison details among three clades and four subgenera. *** $p < 0.001$.

| **Statistic** | **Clade** | | | **Subgenus** | | | |
|---|---|---|---|---|---|---|---|
| | **1** | **2** | **3** | **Amphilaena** | **Eriocoryne** | **Saussurea** | **Theodorea** |
| Number of values | 67 | 50 | 77 | 25 | 30 | 127 | 12 |
| Mean | 0.708 | 0.560 | 0.981 | 0.945 | 0.560 | 0.819 | 0.545 |
| Std. Deviation | 0.452 | 0.121 | 0.385 | 0.434 | 0.160 | 0.428 | 0.069 |
| Std. Error of Mean | 0.055 | 0.017 | 0.044 | 0.087 | 0.029 | 0.038 | 0.020 |
| Lower 95% CI of mean | 0.598 | 0.525 | 0.894 | 0.766 | 0.500 | 0.744 | 0.501 |
| Upper 95% CI of mean | 0.819 | 0.594 | 1.069 | 1.124 | 0.619 | 0.894 | 0.589 |

| **ANOVA summary** | **Clade** | **Subgenus** |
|---|---|---|
| F | 22.1300 | 6.6980 |
| P value | <0.0001 | 0.0003 |
| P value summary | *** | *** |
| R square | 0.1882 | 0.0957 |

**Table S6** Statistic summary of *Saussurea* speciation rates from DR statistic. ANOVA analysis shows the comparison details among three clades and four subgenera. * $p < 0.05$.

| **Statistic** | **Clade** | | | **Subgenus** | | | |
|---|---|---|---|---|---|---|---|
| | **1** | **2** | **3** | **Amphilaena** | **Eriocoryne** | **Saussurea** | **Theodorea** |
| Number of values | 67 | 50 | 77 | 25 | 30 | 127 | 12 |
| Mean | 0.997 | 0.776 | 1.143 | 1.087 | 0.540 | 1.106 | 0.814 |
| Std. Deviation | 1.037 | 0.868 | 0.871 | 1.055 | 0.400 | 1.002 | 0.560 |
| Std. Error of Mean | 0.127 | 0.123 | 0.099 | 0.211 | 0.073 | 0.089 | 0.162 |
| Lower 95% CI of mean | 0.744 | 0.530 | 0.945 | 0.652 | 0.391 | 0.930 | 0.458 |
| Upper 95% CI of mean | 1.250 | 1.023 | 1.340 | 1.523 | 0.690 | 1.282 | 1.169 |

| **ANOVA summary** | **Clade** | **Subgenus** |
|---|---|---|
| F | 2.3460 | 3.2860 |
| P value | 0.0985 | 0.0219 |
| P value summary | - | * |
| R square | 0.0240 | 0.0493 |

**Table S7** Model comparison of HiSSE analysis for four binary traits of *Saussurea*. For each HiSSE model, the maximum negative log-likelihood (Loglik), AIC value corrected for sample size (AICc) and the difference between the best model and each model (ΔAICc) are provided, with the best model denoted in bold. For simplicity, just model number (#) is provided, the corresponding model name is provided in Table S3.

| Model | Stemless vs. cauliferous | | | Glabrous vs. densely haired | | | Absence vs. presence of leafy bracts | | | Capitula solitary vs. numerous | | |
|---|---|---|---|---|---|---|---|---|---|---|---|---|
| (#) | loglik | AICc | Δ AICc | loglik | AICc | Δ AICc | loglik | AIC | Δ AICc | loglik | AIC | Δ AICc |
| 1 | **-354.760** | **741.520** | **0.000** | **-392.311** | **816.623** | **0.000** | **-353.882** | **739.764** | **0.000** | **-391.217** | **814.435** | **0.000** |
| 2 | -367.684 | 747.368 | 5.849 | -411.811 | 835.622 | 19.000 | -374.703 | 761.407 | 21.643 | -403.292 | 818.583 | 4.149 |
| 3 | -369.920 | 749.841 | 8.321 | -410.764 | 831.529 | 14.906 | -365.736 | 741.472 | 1.708 | -407.327 | 824.654 | 10.220 |
| 4 | -380.861 | 771.722 | 30.202 | -411.859 | 833.718 | 17.095 | -376.826 | 763.651 | 23.888 | -419.207 | 848.414 | 33.979 |
| 5 | -388.468 | 784.935 | 43.415 | -411.883 | 831.767 | 15.144 | -388.106 | 784.212 | 44.448 | -419.197 | 846.393 | 31.959 |
| 6 | -378.339 | 766.678 | 25.159 | -411.885 | 833.771 | 17.148 | -385.778 | 781.556 | 41.792 | -419.199 | 848.397 | 33.963 |
| 7 | -382.085 | 772.171 | 30.651 | -411.883 | 831.767 | 15.144 | -389.408 | 786.817 | 47.053 | -419.198 | 846.396 | 31.961 |
| 8 | -384.759 | 787.518 | 45.998 | -405.709 | 829.418 | 12.795 | -377.012 | 772.023 | 32.260 | -406.870 | 831.739 | 17.305 |
| 9 | -391.906 | 795.811 | 54.292 | -410.429 | 832.857 | 16.235 | -442.767 | 897.534 | 157.771 | -409.787 | 831.574 | 17.139 |
| 10 | -372.621 | 763.242 | 21.722 | -399.100 | 816.199 | -0.423 | -364.925 | 747.850 | 8.087 | -404.827 | 827.655 | 13.220 |
| 11 | -377.237 | 766.473 | 24.953 | -411.970 | 835.940 | 19.317 | -379.947 | 771.894 | 32.130 | -407.995 | 827.990 | 13.555 |
| 12 | -377.707 | 765.413 | 23.894 | -409.803 | 829.606 | 12.983 | -382.431 | 774.862 | 35.098 | -419.442 | 848.884 | 34.450 |
| 13 | -384.496 | 776.993 | 35.473 | -411.886 | 831.772 | 15.150 | -392.290 | 792.580 | 52.816 | -419.428 | 846.855 | 32.421 |
| 14 | -373.406 | 760.813 | 19.293 | -406.449 | 826.897 | 10.275 | -375.678 | 765.357 | 25.593 | -409.260 | 832.520 | 18.085 |
| 15 | -381.767 | 773.534 | 32.015 | -411.857 | 833.713 | 17.091 | -388.471 | 786.943 | 47.179 | -419.445 | 848.889 | 34.455 |

| | | | | | | | | | | | | |
|---|---|---|---|---|---|---|---|---|---|---|---|---|
| 16 | -365.948 | 745.895 | 4.376 | -411.847 | 837.694 | 21.072 | -373.621 | 761.242 | 21.479 | -406.713 | 827.427 | 12.992 |
| 17 | -377.418 | 764.836 | 23.316 | -406.300 | 822.599 | 5.977 | -386.383 | 782.765 | 43.001 | -408.151 | 826.302 | 11.868 |
| 18 | -366.018 | 750.036 | 8.517 | -399.085 | 826.170 | 9.547 | -361.735 | 741.470 | 1.706 | -403.654 | 825.308 | 10.873 |
| 19 | -371.980 | 755.961 | 14.441 | -400.975 | 823.949 | 7.326 | -370.758 | 753.516 | 13.752 | -409.894 | 831.788 | 17.354 |
| 20 | -375.023 | 760.046 | 18.527 | -409.734 | 829.468 | 12.845 | -378.992 | 767.985 | 28.221 | -419.692 | 849.383 | 34.949 |
| 21 | -377.918 | 763.836 | 22.316 | -409.381 | 826.762 | 10.140 | -386.868 | 781.736 | 41.972 | -420.047 | 848.095 | 33.660 |
| 22 | -365.505 | 745.010 | 3.491 | -407.581 | 829.162 | 12.540 | -374.943 | 763.887 | 24.123 | -410.085 | 834.170 | 19.735 |
| 23 | -372.746 | 755.491 | 13.972 | -406.792 | 823.584 | 6.961 | -382.974 | 775.948 | 36.185 | -411.287 | 832.574 | 18.139 |
| 24 | -369.801 | 753.602 | 12.082 | -412.781 | 839.562 | 22.940 | -374.165 | 762.330 | 22.566 | -405.495 | 824.991 | 10.556 |
| 25 | -374.349 | 758.697 | 17.177 | -405.655 | 821.309 | 4.687 | -377.959 | 765.917 | 26.153 | -410.004 | 830.008 | 15.573 |

**Table S8** Model comparison of MuSSE analysis for multistate traits of *Saussurea*. ANOVA calculations all preferred models constraining each μ to be equal and allowing λ to vary (free λ) (in bold).

| Traits | Models | Df | lnLik | AIC | Pr(>\|Chi\|) |
|---|---|---|---|---|---|
| Leaf hairs | Null model | 8 | -476.7804 | 969.5609 | NA |
| | Full model | 12 | -465.2930 | 954.5861 | 0.0001 |
| | **Free λ** | **10** | **-465.9750** | **951.9499** | **0.0000** |
| | Free μ | 10 | -472.5959 | 965.1917 | 0.0152 |
| Leaf margin | Null model | 8 | -445.7092 | 907.4184 | NA |
| | Full model | 12 | -437.4504 | 898.9008 | 0.0024 |
| | **Free λ** | **10** | **-437.7320** | **895.4639** | **0.0003** |
| | Free μ | 10 | -442.0879 | 904.1758 | 0.0267 |
| Phyllary rows | Null model | 16 | -496.3758 | 1024.7515 | NA |
| | Full model | 20 | -475.3784 | 990.7568 | 0.0000 |
| | **Free λ** | **18** | **-475.3905** | **986.7810** | **0.0000** |
| | Free μ | 18 | -484.8785 | 1005.7570 | 0.0000 |
| Phyllary types | Null model | 16 | -510.0010 | 1052.0020 | NA |
| | Full model | 20 | -489.7781 | 1019.5562 | 0.0000 |
| | **Free λ** | **18** | **-490.1346** | **1016.2691** | **0.0000** |
| | Free μ | 18 | -494.8785 | 1025.7570 | 0.0000 |

**Table S9** Model comparison of GeoHiSSE analysis for ecological habitants of *Saussurea*. The best fitting model (model 4) is in bold. Detailed explanation of these models are described in Caetano *et al.* (2018).

| Models | Loglik | AIC | AICω |
|---|---|---|---|
| 1 | -477.0073 | 960.0146 | 0.0000 |
| 2 | -475.3456 | 960.6912 | 0.0000 |
| 3 | -464.1829 | 938.3657 | 0.0020 |
| **4** | **-451.9680** | **925.9359** | **0.9980** |

**Table S10** Summary table of ecological factors for correlation with diversification rates of *Saussurea* species.

| **Species** | **ClimatePCA1** | **ClimatiePC2** | **Niche breadth** | **Range size (log-transformed)** |
|---|---|---|---|---|
| *Saussurea alaschanica* | 3.3690 | 1.9563 | 0.7020 | 11.5719 |
| *Saussurea alpina* | 2.7809 | 2.7596 | 0.8106 | 11.8332 |
| *Saussurea americana* | 4.8898 | 1.4700 | 0.4043 | 12.3256 |
| *Saussurea amurensis* | 0.0585 | 2.2897 | 0.9558 | 12.0352 |
| *Saussurea andryaloides* | -0.4562 | 2.4498 | 0.8964 | 12.3186 |
| *Saussurea apus* | -1.6515 | 0.7494 | 0.9872 | 12.2847 |
| *Saussurea baicalensis* | -0.3964 | 2.4470 | 0.9143 | 11.7442 |
| *Saussurea baroniana* | 0.6814 | 1.9771 | 0.5103 | 11.9579 |
| *Saussurea bhutanensis* | 0.8358 | -0.3589 | 0.5518 | 11.2284 |
| *Saussurea brachycephala* | 5.1918 | 1.5862 | 0.4813 | 11.1312 |
| *Saussurea bracteata* | -0.5249 | 2.6505 | 0.6856 | 11.5031 |
| *Saussurea brunneopilosa* | -1.2223 | 0.1659 | 0.7784 | 11.9428 |
| *Saussurea cana* | -0.5162 | 1.4913 | 0.6917 | 11.7411 |
| *Saussurea centiloba* | 0.5460 | -1.8907 | 0.5511 | 11.2188 |
| *Saussurea chabyoungsanica* | 1.0893 | -1.8114 | 0.7209 | 11.6019 |
| *Saussurea chinduensis* | -1.2253 | 0.1629 | 0.6714 | 11.9398 |
| *Saussurea ciliaris* | 0.8592 | -1.6602 | 0.4666 | 11.5514 |
| *Saussurea controversa* | 0.3918 | 2.7634 | 0.8286 | 11.8733 |
| *Saussurea coriolepis* | 0.8537 | -0.9376 | 0.2579 | 11.0487 |
| *Saussurea davurica* | -0.3995 | 2.7568 | 0.7492 | 11.2615 |
| *Saussurea depsangensis* | -1.2577 | 0.8001 | 0.2744 | 11.1145 |
| *Saussurea dulongjiangensis* | 0.7151 | 2.5555 | 0.5066 | 11.6932 |
| *Saussurea elegans* | 0.6263 | 3.0588 | 0.6291 | 11.4987 |
| *Saussurea eriocephala* | 1.0045 | -1.9199 | 0.4025 | 10.6783 |
| *Saussurea eriostemon* | 0.2738 | -2.1589 | 0.6752 | 11.2999 |
| *Saussurea fuscipappa* | -0.0886 | -0.6569 | 0.4196 | 10.8267 |
| *Saussurea globosa* | -0.6434 | -0.6383 | 0.8268 | 11.8351 |
| *Saussurea gossipiphora* | 0.3572 | -2.5494 | 0.1660 | 11.0306 |
| *Saussurea grosseserrata* | 0.8827 | -1.9149 | 0.2771 | 10.9887 |
| *Saussurea henryi* | 3.1980 | 1.7853 | 0.4841 | 10.8122 |

| | | | | |
|---|---|---|---|---|
| *Saussurea hylophila* | 2.3580 | -2.0180 | 0.4466 | 10.7344 |
| *Saussurea integrifolia* | 0.9770 | -0.3291 | 0.6609 | 11.5316 |
| *Saussurea involucrata* | -0.4202 | 2.4858 | 0.4851 | 11.1066 |
| *Saussurea japonica* | 3.1256 | 1.1714 | 0.9633 | 12.6558 |
| *Saussurea kaschgarica* | -0.1071 | 2.8368 | 0.5203 | 10.7650 |
| *Saussurea katochaete* | -0.9714 | -0.4450 | 0.8003 | 11.9048 |
| *Saussurea komaroviana* | 1.1651 | 1.4888 | 0.8619 | 11.2175 |
| *Saussurea kuschakewiczii* | 1.0409 | 3.2574 | 0.6951 | 11.5546 |
| *Saussurea lanata* | -1.4429 | -1.0696 | 0.5035 | 11.2569 |
| *Saussurea langpoensis* | -0.6029 | 2.5725 | 0.5376 | 11.3109 |
| *Saussurea laniceps* | -0.4699 | -1.1190 | 0.6513 | 11.7402 |
| *Saussurea latifolia* | 0.7510 | 2.6599 | 0.6679 | 11.5941 |
| *Saussurea leucophylla* | -0.8810 | 2.5626 | 0.6970 | 11.5576 |
| *Saussurea licentiana* | 2.5057 | 1.2372 | 0.5725 | 11.5720 |
| *Saussurea longifolia* | -0.1610 | -0.9535 | 0.7735 | 12.0006 |
| *Saussurea malitiosa* | -1.8243 | 0.9901 | 0.2051 | 10.8015 |
| *Saussurea merinoi* | 3.1402 | 1.6185 | 0.6089 | 11.3554 |
| *Saussurea mucronulata* | 0.5538 | 2.9254 | 0.9141 | 12.0353 |
| *Saussurea muliensis* | 1.3279 | -0.6061 | 0.7104 | 11.4264 |
| *Saussurea nigrescens* | 0.0178 | 0.4702 | 0.7412 | 11.8343 |
| *Saussurea nuda* | 0.7586 | 2.4155 | 0.5930 | 11.6002 |
| *Saussurea obvallata* | 0.4758 | -0.7189 | 0.7541 | 11.8451 |
| *Saussurea odontolepis* | 0.8036 | 2.2819 | 0.8929 | 11.5209 |
| *Saussurea orgaadayi* | -1.2104 | 2.5484 | 0.0482 | 10.6942 |
| *Saussurea pachyneura* | 0.6164 | -1.0334 | 0.9984 | 11.9876 |
| *Saussurea paucijuga* | -1.6463 | 1.1593 | 0.9842 | 12.0784 |
| *Saussurea peduncularis* | 0.8133 | -2.0874 | 0.2002 | 10.8398 |
| *Saussurea petrovii* | 0.5266 | 1.9191 | 0.6850 | 11.5449 |
| *Saussurea picridifolia* | 1.0333 | -1.3390 | 0.1708 | 10.7200 |
| *Saussurea pinnatidentata* | -0.7834 | 2.3758 | 0.7194 | 11.4561 |
| *Saussurea polylepis* | 3.9680 | 2.4637 | 0.9577 | 10.7321 |
| *Saussurea poochlamys* | 0.8243 | -1.4399 | 0.6298 | 11.3638 |
| *Saussurea populifolia* | 1.4534 | 0.1048 | 0.7947 | 11.5056 |
| *Saussurea pseudoalpina* | -1.0098 | 2.5770 | 0.6609 | 11.1444 |

| | | | | |
|---|---|---|---|---|
| *Saussurea pseudosimpsoniana* | -1.8795 | 0.0187 | 0.5413 | 10.6870 |
| *Saussurea pseudotridactyla* | -1.5516 | -0.5851 | 0.1865 | 11.3949 |
| *Saussurea pulchella* | 0.0176 | 1.9988 | 0.8636 | 11.7812 |
| *Saussurea pulvinata* | -2.1234 | 1.4481 | 0.2792 | 11.0705 |
| *Saussurea romuleifolia* | 0.1513 | -1.4357 | 0.6463 | 11.5542 |
| *Saussurea runcinata* | -1.0114 | 2.1478 | 0.5085 | 11.2281 |
| *Saussurea salicifolia* | -0.1126 | 1.9669 | 0.8008 | 12.0792 |
| *Saussurea salsa* | 0.6862 | 3.2972 | 0.8559 | 12.0926 |
| *Saussurea semiamplexicaulis* | 0.4710 | -0.3726 | 0.6339 | 11.4639 |
| *Saussurea semifasciata* | -0.0122 | -0.8049 | 0.8023 | 11.6154 |
| *Saussurea semilyrata* | 0.0335 | -1.4360 | 0.7361 | 11.5081 |
| *Saussurea simpsoniana* | 0.0147 | -1.0309 | 0.7518 | 11.6335 |
| *Saussurea sobarocephala* | 0.2644 | 0.1653 | 0.5404 | 11.3957 |
| *Saussurea spathulifolia* | -0.1270 | -1.5581 | 0.3878 | 11.0720 |
| *Saussurea stella* | -0.3924 | -0.5540 | 0.9690 | 12.2351 |
| *Saussurea stricta* | 1.5591 | 1.2228 | 0.8026 | 11.5281 |
| *Saussurea subtriangulata* | 2.1158 | 2.8442 | 0.3884 | 11.1346 |
| *Saussurea subulata* | -1.7602 | 1.6421 | 0.9833 | 12.0855 |
| *Saussurea superba* | -1.0759 | -0.1699 | 0.7953 | 11.7667 |
| *Saussurea sutchuenensis* | 1.9874 | 0.6827 | 0.5030 | 10.7043 |
| *Saussurea tangutica* | -1.6735 | -0.0621 | 0.4018 | 11.4716 |
| *Saussurea thomsonii* | -2.0415 | 1.6225 | 0.7953 | 11.8603 |
| *Saussurea thoroldii* | -2.1023 | 0.7033 | 0.6051 | 11.6224 |
| *Saussurea tianshuiensis* | 1.0863 | -1.8144 | 0.2986 | 11.1128 |
| *Saussurea tridactyla* | -1.2285 | -1.1503 | 0.8021 | 10.9055 |
| *Saussurea tunglingensis* | 0.3239 | 1.1162 | 0.4386 | 11.7122 |
| *Saussurea uliginosa* | 0.5900 | -1.8838 | 0.3902 | 11.2870 |
| *Saussurea uniflora* | 0.5273 | -2.3574 | 0.5226 | 10.9451 |
| *Saussurea veitchiana* | 3.7643 | 1.4672 | 0.3643 | 11.2885 |
| *Saussurea velutina* | -0.1549 | -1.0014 | 0.7437 | 11.2207 |
| *Saussurea wellbyi* | -2.0311 | 0.1355 | 0.6946 | 11.1528 |
| *Saussurea woodiana* | 0.2775 | -0.2582 | 0.9953 | 11.9165 |
| *Saussurea hookeri* | -1.1419 | -0.7870 | 0.7323 | 11.3681 |
| *Saussurea obvallata* | 0.4788 | -0.7159 | 0.7560 | 11.7031 |

| | | | | |
|---|---|---|---|---|
| *Saussurea pubifolia* | -0.9590 | -0.6616 | 0.2748 | 12.2441 |
| *Saussurea gossipiphora* | 0.4502 | -2.4564 | 0.2245 | 10.8075 |
| *Saussurea tridactyla* | -1.3725 | -1.2943 | 0.7114 | 11.3503 |
| *Saussurea gnaphalodes* | -1.3882 | 1.7144 | 0.6155 | 11.1266 |
| *Saussurea salwinensis* | -0.1195 | -0.7957 | 0.6057 | 11.6462 |
| *Saussurea przewalskii* | -0.5151 | -0.7307 | 0.9884 | 12.3276 |
| *Saussurea delavayi* | 1.2081 | -2.5444 | 0.3409 | 10.9964 |
| *Saussurea leontodontoides* | -0.8134 | -0.0490 | 0.9182 | 12.0234 |
| *Saussurea kingii* | -1.6857 | -1.0230 | 0.3821 | 10.4244 |
| *Saussurea japonica* | 3.0176 | 1.0634 | 0.9519 | 12.3464 |
| *Saussurea_acutisquama* | -0.9020 | -0.4353 | 0.4833 | 10.7203 |
| *Saussurea_aster* | -1.8954 | -0.1402 | 0.7207 | 12.2448 |
| *Saussurea_bodinieri* | 0.2843 | -0.7626 | 0.9608 | 11.4066 |
| *Saussurea_caudata* | 0.0976 | -0.9164 | 0.7842 | 11.7545 |
| *Saussurea_cauloptera* | 3.7498 | 1.8179 | 0.9885 | 11.8972 |
| *Saussurea_ceterach* | -0.9891 | -0.6712 | 0.8291 | 11.5749 |
| *Saussurea_cochlearifolia* | 1.3303 | -1.0420 | 0.3661 | 11.7687 |
| *Saussurea_columnaris* | 0.2924 | -1.4684 | 0.8099 | 10.7950 |
| *Saussurea_compta* | 1.2121 | -1.0256 | 0.6375 | 11.1704 |
| *Saussurea_conica* | -0.8360 | -1.0434 | 0.0229 | 11.2464 |
| *Saussurea_dolichopoda* | 3.3112 | 1.7895 | 0.8772 | 11.1727 |
| *Saussurea_dzeurensis* | -1.1753 | -0.5101 | 0.4743 | 11.3104 |
| *Saussurea_epilinophylla* | -1.6176 | -0.6511 | 0.4994 | 11.4253 |
| *Saussurea_erubescens* | -0.9606 | 0.0023 | 0.9089 | 11.3907 |
| *Saussurea_georgei* | -0.5051 | -1.0301 | 0.5504 | 11.6033 |
| *Saussurea_globosa* | -0.5324 | -0.5273 | 0.8967 | 11.8594 |
| *Saussurea_graminea* | -1.0364 | -0.6397 | 0.8408 | 11.2500 |
| *Saussurea_graminifolia* | 0.6373 | -0.7902 | 0.6648 | 11.8321 |
| *Saussurea_hieracioides* | -0.7785 | -1.0982 | 0.7276 | 11.9944 |
| *Saussurea_hypsipeta* | -1.8825 | 0.0157 | 0.5252 | 11.3235 |
| *Saussurea_kansuensis* | -0.7140 | -0.6604 | 0.8707 | 12.3114 |
| *Saussurea_katochaete* | -0.9054 | -0.3790 | 0.8387 | 11.1313 |
| *Saussurea_kungii* | 2.5301 | 1.8590 | 0.6085 | 11.4746 |
| *Saussurea_lavrenkoana* | -0.3351 | -1.6346 | 0.5671 | 12.0031 |

| | | | | |
|---|---|---|---|---|
| *Saussurea_leclerei* | 0.5681 | -1.6638 | 0.7002 | 11.3329 |
| *Saussurea_leiocarpa* | -0.1289 | 0.5405 | 0.7423 | 10.9637 |
| *Saussurea_leucoma* | 0.2096 | -1.5820 | 0.7846 | 10.9430 |
| *Saussurea_likiangensis* | 0.2384 | -0.6756 | 0.8544 | 11.1940 |
| *Saussurea_loriformis* | -0.3213 | -0.8904 | 0.2638 | 11.5057 |
| *Saussurea_luae* | -0.8725 | -0.9761 | 0.5132 | 11.8032 |
| *Saussurea_macrota* | 0.9453 | 0.7126 | 0.8543 | 10.9945 |
| *Saussurea_medusa* | -1.1273 | -0.7013 | 0.5544 | 10.6675 |
| *Saussurea_nyalamensis* | -0.4406 | -0.4692 | 0.4004 | 11.3585 |
| *Saussurea_oligantha* | 2.7862 | 1.5083 | 0.6429 | 12.1385 |
| *Saussurea_pachyneura* | 0.4754 | -1.1744 | 0.9396 | 11.1325 |
| *Saussurea_paxiana* | -1.2271 | -0.0742 | 0.6854 | 11.5816 |
| *Saussurea_phaeantha* | -1.0432 | 0.0293 | 0.7553 | 11.9606 |
| *Saussurea_pilobioides* | 3.7408 | 1.8089 | 0.7432 | 11.7773 |
| *Saussurea_polycolea* | -0.4704 | -0.5095 | 0.8717 | 11.6390 |
| *Saussurea_pseudosimpsoniana* | -1.6665 | 0.2317 | 0.8225 | 11.7353 |
| *Saussurea_pygmaea* | 3.4087 | 1.2249 | 0.0739 | 10.9619 |
| *Saussurea_quercifolia* | -0.2670 | -1.0069 | 0.6373 | 11.5732 |
| *Saussurea_recurvata* | 0.1663 | 2.5519 | 0.6167 | 11.0731 |
| *Saussurea_scabrida* | 0.4091 | -0.8910 | 0.7946 | 11.6047 |
| *Saussurea_sericea* | -2.3114 | -0.4326 | 0.1873 | 11.1592 |
| *Saussurea_spathulifolia* | -0.0820 | -1.5131 | 0.8022 | 11.3278 |
| *Saussurea_spatulifolia* | 0.1168 | -1.6163 | 0.3726 | 11.0480 |
| *Saussurea_stella* | -0.7914 | -0.9530 | 0.7176 | 11.8361 |
| *Saussurea_stoliczkae* | -0.9100 | -0.3088 | 0.8273 | 11.8820 |
| *Saussurea_subulata* | -1.8982 | 1.5041 | 0.8964 | 11.9475 |
| *Saussurea_subulisquama* | -0.9961 | 0.0648 | 0.8818 | 11.8512 |
| *Saussurea_sylvatica* | -1.0023 | -0.2579 | 0.6560 | 11.4615 |
| *Saussurea_velutina* | -0.0889 | -0.9354 | 0.7853 | 11.4168 |
| *Saussurea_wardii* | 0.1446 | -1.2735 | 0.8198 | 11.5487 |
| *Saussurea_wernerioides* | -1.2262 | -0.8094 | 0.7654 | 11.6929 |

**Table S11** Model comparison of QuaSSE analysis for correlation between ecological factors and speciation rates of *Saussurea,* with the best model shown in bold.

| ClimatePC1 | | | | | |
|---|---|---|---|---|---|
| | **Df** | **lnLik** | **AIC** | **ChiSq** | **Pr(>\|Chi\|)** |
| Constant | 3 | -620.254 | 1246.508 | *NA* | *NA* |
| **Linear** | **4** | **-616.274** | **1240.548** | **7.960** | **0.005** |
| Sigmoidal | 6 | -614.881 | 1241.762 | 10.746 | 0.013 |
| Drift + linear | 5 | -616.102 | 1242.203 | 8.305 | 0.016 |
| Drift + sigmoidal | 7 | -614.226 | 1242.453 | 12.056 | 0.017 |
| ClimatePC2 | | | | | |
| **Constant** | **3** | **-588.762** | **1183.524** | ***NA*** | ***NA*** |
| Linear | 4 | -588.760 | 1185.520 | 0.003 | 0.953 |
| Sigmoidal | 6 | -587.975 | 1187.951 | 1.573 | 0.666 |
| Drift + linear | 5 | -588.675 | 1187.350 | 0.174 | 0.917 |
| Drift + sigmoidal | 7 | -587.718 | 1189.437 | 2.087 | 0.720 |
| Niche breadth | | | | | |
| Constant | 3 | -275.741 | 557.482 | *NA* | *NA* |
| Linear | 4 | -275.741 | 559.482 | 0.000 | 1.000 |
| Sigmoidal | 6 | -264.119 | 540.238 | 23.243 | 0.000 |
| Drift + linear | 5 | -261.221 | 532.441 | 29.041 | 0.000 |
| **Drift + sigmoidal** | **7** | **-257.766** | **529.532** | **35.950** | **0.000** |
| Species rang size | | | | | |
| Constant | 3 | -359.810 | 725.621 | *NA* | *NA* |
| Linear | 4 | -359.810 | 727.621 | 0.000 | 1.000 |
| Sigmoidal | 6 | -347.704 | 707.408 | 24.213 | 0.000 |
| Drift + linear | 5 | -359.810 | 729.620 | 0.000 | 1.000 |
| **Drift + sigmoidal** | **7** | **-343.336** | **700.671** | **32.949** | **0.000** |

### Reference:

**Beaulieu JM, O'Meara BC. 2016.** Detecting Hidden Diversification Shifts in Models of Trait-Dependent Speciation and Extinction. *Systematic Biology* **65**(4): 583-601.

**Caetano DS, O'Meara BC, Beaulieu JM. 2018.** Hidden state models improve state-dependent diversification approaches, including biogeographical models. *Evolution* **72**(11): 2308-2324.

**Jetz W, Thomas GH, Joy JB, Hartmann K, Mooers AO. 2012.** The global diversity of birds in space and time. *Nature* **491**(7424): 444-448.
